# Light induced synaptic vesicle autophagy

**DOI:** 10.1101/440719

**Authors:** Sheila Hoffmann, Marta Orlando, Ewa Andrzejak, Thorsten Trimbuch, Christian Rosenmund, Frauke Ackermann, Craig C. Garner

## Abstract

The regulated turnover of synaptic vesicle (SV) proteins is thought to involve the ubiquitin dependent tagging and degradation through endo-lysosomal and autophagy pathways. Yet, it remains unclear which of these pathways are used, when they become activated and whether SVs are cleared en-mass together with SV proteins or whether both are degraded selectively. Equally puzzling is how quickly these systems can be activated and whether they function in real time to support synaptic health. To address these questions, we have developed an imaging based system that simultaneously tags presynaptic proteins while monitoring autophagy. Moreover, by tagging SV proteins with a light activated reactive oxygen species (ROS) generator, Supernova, it was possible to temporally control the damage to specific SV proteins and assess their consequence to autophagy mediated clearance mechanisms and synaptic function. Our results show that, in mouse hippocampal neurons, presynaptic autophagy can be induced in as little as 5-10 minutes and eliminates primarily the damaged protein rather than the SV en-mass. Importantly, we also find that autophagy is essential for synaptic function, as light-induced damage to e.g. Synaptophysin only compromises synaptic function when autophagy is simultaneously blocked. These data support the concept that presynaptic boutons have a robust highly regulated clearance system to maintain not only synapse integrity, but also synaptic function.

**Significance Statement:** The real-time surveillance and clearance of synaptic proteins is thought to be vital to the health, functionality and integrity of vertebrate synapses and is compromised in neurodegenerative disorders, yet the fundamental mechanisms regulating these systems remain enigmatic. Our analysis reveals that presynaptic autophagy is a critical part of a real-time clearance system at glutamatergic synapses capable of responding to local damage of synaptic vesicle proteins within minutes and to be critical for the ongoing functionality of these synapses. These data indicate that synapse autophagy is not only locally regulated but also crucial for the health and functionality of vertebrate presynaptic boutons.

## Introduction

The integrity of vertebrate synapses requires robust cellular programs that monitor the activity states of thousands of proteins, eliminating those that are mis-folded or damaged. Failure of these programs can lead to the accumulation of non-functional proteins that reduce the efficiency of synaptic transmission and promote neurodegeneration (Liang and Sigrist, 2018; Vijayan and Verstreken, 2017; Waites et al., 2013). Neurons are endowed with several surveillance and clearance systems. These include an ubiquitin based tagging system that conjugates ubiquitin chains to damaged proteins, as well as several degradative systems that, for example, eliminate soluble proteins via the proteasome or integral membrane proteins and protein aggregates via the endo-lysosomal and/or autophagy systems (Wang et al., 2017).

Given their distance from the cell soma and high metabolic demand, synapses poise a significant challenge to neurons, as they have to maintain and ensure a stable functional pool of proteins (Tammineni et al., 2017). How might this be achieved? Emerging data indicate that synapses utilize their own local machinery to eliminate proteins e.g. in response to changes in synaptic activity or homeostatic plasticity (Vijayan and Verstreken, 2017). For example, the ESCRT system facilitates via Rab35 the elimination of subsets of SV proteins in response to changes in synaptic activity (Sheehan et al., 2016). Moreover, specific E3 ubiquitin ligases have been associated with the selective removal of key regulators of synaptic transmission such as RIM1 (Yao et al., 2007) and Munc13 by the proteasome (Jiang et al., 2010; Yi and Ehlers, 2005). Intriguingly, two active zone proteins, Piccolo and Bassoon, have also been identified as regulators of presynaptic proteostasis, as their inactivation leads to the loss of SVs and disintegration of synaptic junctions through the activation of E3 ligases (Waites et al., 2013) and autophagy (Okerlund et al., 2017).

Although these clearance systems are anticipated to ensure functionality of synaptic proteins, it remains unclear whether some are specialized for the removal of only subsets of synaptic proteins. A growing number of studies point to the importance of macroautophagy not only in maintaining mitochondrial health, but also the clearance of aggregated proteins (Vijayan and Verstreken, 2017). Interestingly, in Alzheimer’s disease brains, an up-regulation of autophagy has been observed (Boland et al., 2008; Lee et al., 2010; Nixon et al., 2005), however, in other diseases characterized by aggregate-prone proteins such as Parkinson’s and Huntington’s disease, autophagy is not engaged (Martinez-Vicente et al., 2010; Nixon, 2013; Rubinsztein et al., 2012; Spencer et al., 2009), which might contribute to the accumulation of protein aggregates and subsequent reduced neuronal survival (Ebrahimi-Fakhari et al., 2011; Nixon, 2013; Yue et al., 2009). This latter concept is supported by the analysis of Atg5 or Atg7 knockout mice, two essential autophagy-related proteins, which exhibit hallmarks of neurodegeneration (Hara et al., 2006; Komatsu et al., 2006). Defining the role of degradative systems during health and disease requires a better understanding of when and where each is turned on and which subsets of proteins they eliminate. For example, those critical for the real-time maintenance of synaptic function should be locally regulated and operating on a second to minute time scale, while those responding to chronic damage may act on longer time scales like hours. To address these fundamental questions, we have developed a strategy to selectively damage SV proteins within presynaptic boutons. This was accomplished by tethering the light activated free radical oxygen species (ROS) generator Supernova (Takemoto et al., 2013) to different SV proteins, allowing the local light induced damage of SV proteins with a half-radius of photo-damage as small as 3-4nm (Takemoto et al., 2013).

This manipulation was found to rapidly and selectively induce presynaptic autophagy within 5 minutes and lead primarily to the elimination of damaged proteins and not SV proteins en-mass. Moreover, the selective damage of SV proteins allowed us to show that presynaptic autophagy is critical for the real-time maintenance of synaptic transmission.

## Material and Methods

### Construction of vectors

Monitoring of autophagy within presynaptic boutons was achieved by creating a set of lentiviral expression vectors. All vectors are based on the commercially available vector FUGW (Addgene). In order to co-express mCherry-tagged Synaptophysin (Syp) and eGFP-LC3, Synaptophysin-mCherry (Synaptophysin, NM_012664.3) was synthesized by Eurofins Genomics with a downstream glycine linker that was fused to a self-cleaving 2A peptide (Kim et al., 2011). This element was then exchanged with GFP in the FUGW vector by ligation. Subsequently, the eGFP-LC3 (LC3, U05784.1) segment from FU-ptf-LC3 (Okerlund et al., 2017) was subcloned in frame after the P2A sequence, which resulted in the vector FU-Syp-mCherry-P2A-eGFP-LC3. This vector also served as a template for tagging Synaptophysin with Supernova. Here, Supernova was synthesized by Eurofins Genomics (Supernova, AB522905) (Takemoto et al., 2013) and exchanged for mCherry forming FU-Syp-Supernova-P2A-eGFP-LC3. To monitor endolysosomal systems PCR amplified Rab7 (XM_005632015.2) was exchanged with LC3 in FU-Syp-Supernova-P2A-eGFP-LC3 by Gibson assembly (Gibson et al., 2009) creating FU-Syp-Supernova-P2A-eGFP-Rab7. Lentiviral vectors expressing eGFP-LC3 and either Supernova tagged Synapsin (Syn) (NM_019133) or Synaptotagmin (Syt) (NM_001252341) (FU-Syn-Supernova-P2A-eGFP-LC3 and FU-Syt-Supernova-P2A-eGFP-LC3) were created by PCR amplification of Synapsin or Synaptotagmin from plasmid DNA (Chang et al., 2018; Waites et al., 2013) before being subjected to a Gibson assembly reaction with the purified Syp-deleted FU-Syp-Supernova-P2A-eGFP-LC3 vector. All final constructs were verified by both restriction digest and sequencing.

### HeLa cell culture and infection

HeLa cells were maintained in DMEM complete medium (DMEM, 10%FCS, 1%P/S) (Thermo Fisher Scientific, Waltham, USA). Medium was changed every 2 to 4 days. HeLa cells were routinely passaged at 80% confluence. Cells were washed with PBS and subsequently treated with 0.05% Trypsin-EDTA (Thermo Fisher Scientific, Waltham, USA) for 1 min at 37°C. Trypsin was inhibited using DMEM complete medium, afterwards cells were detached from the flask, counted and re-plated at a density of 30k per 1cm^2^ onto glass coverslips. 24 hours after plating, HeLa cells were infected with lentivirus adding 100μl per 6-well. 3 days after infection, DMEM was exchanged to EBSS medium (Thermo Fisher Scientific, Waltham, USA) containing 100μM chloroquine (Sigma-Aldrich, St. Louis, USA) for 2 hours at 37°C, in order to enhance the visualization of autophagy by blocking lysosomal degradation. Control cells were left untreated.

### Immunocytochemistry of HeLa cells

Cells were fixed with 4% PFA in PBS for 4 min at RT and washed with PBS twice. All following steps were performed at RT. Cells were permeabilized by three washing steps with PBS + 0.2% Tween-20 (PBS-T) for a total of 30 min followed by incubation with PBS-T with 5% normal goat serum (NGS) (=blocking solution) for another 30 min. The primary antibody was diluted in blocking solution and cells were incubated in this solution for 45 min. The following antibodies were used: primary antibodies against LC3 (1:500; rabbit; MBL International, Woburn, USA; Cat# PM036Y), p62 (1:200; mouse; BD, Heidelberg, Germany; Cat# 610833). Afterwards cells were washed three times in PBS-T for 10 min each. The secondary antibody, diluted in PBS-T 1:1000 (Thermo Fisher Scientific, Waltham, USA), was put onto the cells for 60 min and washed away twice with PBS-T and once with PBS for 10 min each. Finally, coverslips were mounted using ProLong Diamond Antifade Mountant (Thermo Fisher Scientific, Waltham, USA).

### Preparation of cultured hippocampal neurons

All procedures for experiments involving animals were approved by the animal welfare committee of Charité Medical University and the Berlin state government. For live cell imaging and immunocytochemistry, hippocampal neuron cultures were prepared on glass coverslips using the Banker protocol (Banker, 1988; Meberg and Miller, 2003) or on μ-Slide 8 Well culture dishes (ibidi GmbH, Martinsried, Germany). For the first, astrocytes from mouse WT cortices P0-2 were seeded on 6-well or 12-well plates at a density of 10k per 1cm^2^ 5-7 d before neuron preparation. Then, hippocampi were dissected from WT mice P0-2 brains in cold Hanks’ Salt Solution (Millipore, Darmstadt, Germany), followed by a 30 min incubation in enzyme solution (DMEM (Gibco, Thermo Fisher Scientific, Waltham, USA), 3.3mM Cystein, 2mM CaCl_2_, 1mM EDTA, 20U/ml Papain (Worthington, Lakewood, USA)) at 37°C. Papain reaction was inhibited by the incubation of hippocampi in inhibitor solution DMEM, 10% fetal calf serum (FCS) (Thermo Fisher Scientific, Waltham, USA), 38mM BSA (Sigma-Aldrich, St. Louis, USA) and 95mM Trypsin Inhibitor (Sigma-Aldrich, St. Louis, USA) for 5 min. Afterwards, cells were triturated in NBA (Neurobasal-A Medium, 2% B27, 1% Glutamax, 0.2%P/S) (Thermo Fisher Scientific, Waltham, USA) by gentle pipetting up and down. Isolated cells were plated onto nitric acid washed and poly-l-lysine coated glass coverslips with paraffin dots at a density of 10k per 1cm^2^. After 1.5 hours the coverslips were put upside down onto the prepared astrocytes and co-cultured in NBA at 37°C, 5% CO_2_, for 13-15 d (days in vitro, DIV) before starting experiments. For the second, dissociated hippocampal neurons were plated directly onto μ-Slide 8 Well Grid-500 ibiTreat culture dishes (ibidi GmbH, Martinsried, Germany) at a density of 25k per 1cm^2^ and maintained in NBA at 37°C, 5% CO_2_, for 13-15 d before starting experiments.

### Lentivirus production

All lentiviral particles were provided by the Viral Core Facility of the Charité - Universitätsmedizin Berlin (vcf.charite.de) and were prepared as described previously. Briefly, HEK293T cells were cotransfected with 10μg of shuttle vector, 5μg of helper plasmid pCMVdR8.9, and 5μg of pVSV.G with X-tremeGENE 9 DNA transfection reagent (Roche Diagnostics, Mannheim, Germany). Virus containing cell culture supernatant was collected after 72 hours and filtered for purification. Aliquots were flash-frozen in liquid nitrogen and stored at −80°C.

### Immunocytochemistry of hippocampal neurons

Primary hippocampal neurons (expressing FU-Syp-mCherry-P2A-eGFP-LC3), 13-15 DIV, were treated with 2μM rapamycin (Sigma-Aldrich, St. Louis, USA) for either 10 min or 2h, or for the 10 min time point with 1μM wortmannin (InvivoGen, San Diego, USA) additionally. Untreated cells were used as a control. After treatment, cells were fixed with 4% PFA in PBS for 4 min and washed twice with PBS (10 min each). Afterwards, cells were permeabilized with PBS + 0.2% Tween-20 (PBS-T) three times for 10 min each. Following a 30 min incubation with 5% normal goat serum (NGS) in PBS-T (=blocking solution), neurons were incubated with primary antibodies, diluted in blocking solution, for 45 min at RT. The following antibodies were used: primary antibody against p62 (1:500; rabbit; MBL International, Woburn, USA; Cat# PM045), Homer1 (1:1000; guinea pig; synaptic systems; Göttingen, Germany; Cat# 160004), Killerred (recognizes Supernova) (1:1000; rabbit; evrogen, Moscow, Russia; Cat# AB961), GFP (1:1000; chicken; Thermo scientific, Waltham, USA; Cat# A10262), Bassoon (1:500; guinea pig; synaptic systems, Göttingen, Germany; Cat# 141004), Synaptotagmin1 (1:1000; mouse; synaptic systems, Göttingen, Germany; Cat# 105011), Synaptophysin1 (1:1000; mouse; synaptic systems, Göttingen, Germany; Cat# 101011), Synapsin1 (1:1000; rabbit; abcam, Cambridge, UK; Cat# ab64581), Chmp2b (1:200; rabbit; abcam, Cambridge, UK; Cat# ab33174). Afterwards cells were washed three times in PBS-T for 10 min each, incubated with the secondary antibody, diluted in PBS-T 1:1000 (Thermo Fisher Scientific, Waltham, USA), for 60 min and washed twice with PBS-T and once with PBS for 10 min each. Finally, coverslips were dipped in H_2_O and mounted in ProLong Diamond Antifade Mountant (Thermo Fisher Scientific, Waltham, USA).

### Western Blot analyses

Cultured hippocampal neurons, either infected with lentivirus at 2-3 DIV (TD) or uninfected (UT), were grown on 6-well-plates with a density of 20k per 1cm^2^ until 13-15 DIV. All following steps were performed at 4°C. Neurons were kept on ice and washed twice with cold PBS. Subsequently, cells were detached by mechanical force. Isolated cells were centrifuged at 4000rpm for 10 min and resuspended in 100μl lysis buffer (50mM Tris pH 7.9, 150mM NaCl, 5mM EDTA, 1% Triton X-100, 1% NP-40, 0.5% Deoxycholate, protease inhibitor cOmplete Tablets 1x) and incubated for 5 min on ice. Afterwards, cell suspension was centrifuged at 13000rpm for 10 min after which the supernatant was transferred into a new tube. Subsequently, the protein concentration was determined using the Pierce BCA Protein Assay Kit (Thermo Fisher Scientific, Waltham, USA). The same amount of total protein was then separated by SDS-PAGE and transferred onto a PVDF membrane. Afterwards, the membrane was blocked in 5% milk in TBS-T (20mM Tris, 150mM NaCl, 0.1% Tween-20) for 1 hour followed by primary antibody incubation (1:1000 in 3% milk in TBS-T) over night at 4°C. The following antibodies were used: primary antibody against mCherry (1:1000; rabbit; abcam, Cambridge, UK; Cat# ab167453). Afterwards, the membrane was washed three times with TBS-T for 10 min each and incubated with the secondary antibody (1:2500 in 3% milk in TBS-T) for 1 hour at RT. HRP-conjugated secondary antibodies were diluted 1:25000 (Sigma-Aldrich, St. Louis, USA). Afterwards, the membrane was washed three times with TBS-T and bands were visualized using 20x LumiGLO Reagent and 20x Peroxidase (Cell Signaling, Danvers, USA).

### Photo-bleaching primary hippocampal neurons expressing Supernova-constructs

Primary hippocampal neurons in μ-Slide 8 Well Grid-500 ibiTreat (ibidi GmbH, Martinsried, Germany) culture dishes expressing Syp-Supernova, Syn-Supernova or Syt-Supernova cassettes were imaged at 13-15 DIV in Neurobasal Medium without phenol red (Thermo Fisher Scientific, Waltham, USA) at 37°C. Afterwards, a smaller diaphragm restricted area within the field of view was bleached for 60 seconds using 581nm wavelength light from a mercury lamp (100% HXP 120 V, 43 HE filter set 563/581). Immediately after bleaching, a second image was taken confirming the radius of the bleached area. Neurons were fixed at different time points (2-10 min, 56-64 min, 116-124 min) after bleaching Supernova and immunostained with antibodies against Supernova (using a Killerred antibody), GFP, Bassoon or Chmp2b (for procedure see immunocytochemistry of hippocampal neurons). For autophagy inhibition, 1μM wortmannin was added right before the bleaching and kept on the cells till they were fixed. To trigger the dispersion of Synapsin-Supernova before bleaching, fields of views were imaged, the medium was changed to tyrodes buffer 60mM KCl, followed by immediate bleaching for 60 seconds. Subsequently, tyrodes buffer 60mM KCl was exchanged with Neurobasal medium without phenol red and images were taken. Afterwards, neurons were returned to a 37C incubator and fixed after 1 hour. After immunostaining, the same fields of view including the bleached areas were imaged utilizing the grid on the μ-Slide 8 Well Grid-500 culture dishes.

### Basal autophagy in primary hippocampal neurons

Primary hippocampal neurons in μ-Slide 8 Well Grid-500 ibiTreat (ibidi GmbH, Martinsried, Germany) culture dishes expressing FU-eGFP-LC3 were left untreated and fixed at 13-15 DIV. Afterwards, neurons were immunostained with antibodies against GFP, Bassoon and Synaptophysin1/Synapsin1/Synaptotagmin1 (for procedure see immunocytochemistry of hippocampal neurons).

### FM dye uptake

Primary hippocampal neurons, expressing FU-Syp-Supernova-P2A-eGFP-LC3 or FU-Syn-Supernova-P2A-eGFP-LC3, were used for live cell experiments. These were performed using a custom-built imaging chamber designed for liquid perfusion at 37°C. Cells were imaged in tyrodes buffer pH 7.4 (119mM NaCl, 2.5mM KCl, 25mM HEPES, 2mM CaCl_2_, 2mM MgCl_2_, 30mM glucose) and stimulated for 90s in 90mM KCl buffer (31.5mM NaCl, 90mM KCl, 25mM HEPES, 2mM CaCl_2_, 2mM MgCl_2_, 30mM glucose) containing FM 1-43 dye (Thermo Fisher Scientific, Waltham, USA) at a final concentration of 1μg per ml. After stimulation, cells were washed with 20ml tyrodes buffer and subsequently imaged. To inhibit autophagy, 1μM wortmannin was added to neurons 1 min before light activation of Supernova and ~5 min before stimulation. Note, wortmannin was present in all solutions (90mM KCl FM dye, tyrodes buffer washing).

### Electron microscopy

Cultured hippocampal neurons were plated on astrocytes on 6mm sapphire disks at a density of 20k per 1cm^2^ and infected with FU-Syp-Supernova-P2A-eGFP-LC3 at 2-3 DIV. To better correlate regions of interest at the fluorescence and electron microscopy level, carbon was coated in the shape of an alphabetical grid on sapphire disks with the help of a metal mask (finder grid, Plano GmbH, Wetzlar, Germany). After a total of 13-15 days in culture, the sapphire disks were transferred into uncoated μ-Slide 8 Well to perform the bleaching experiment (for procedure see bleaching of primary hippocampal neurons expressing Supernova-constructs). Cryo-fixation using a high pressure freezing machine (EM-ICE, Leica, Wetzlar, Germany) was conducted at different time points after bleaching (10 min, 40 min) in Neurobasal medium without phenol red with the addition of a drop 10% Ficoll solution (Sigma-Aldrich, St. Louis, USA) to prevent ice crystal damage. After freezing, samples were cryo-substituted in anhydrous acetone containing 1% glutaraldehyde, 1% osmium tetroxide and 1% milliQ water in an automated freeze-substiution device (AFS2, Leica). The temperature was kept for 4 hours at -90°C, brought to -20°C (5°C/h), kept for 12 hours at -20°C and then brought from -20°C to +20°C. Once at room temperature, samples were *en-bloc* stained in 0.1% uranyl acetate, infiltrated in increasing concentration of Epoxy resin (Epon 812, EMS Adhesives, Delaware, USA) in acetone and finally embedded in Epon for 48 hours at 65°C. Sapphire disks were removed from the cured resin block by thermal shock. At this point the alphabetical grid was visible on the resin block and was used to find back the bleached regions. The corresponding areas were excised from the blocks for ultrathin sectioning. For each sapphire, as a control, an additional resin blocks was excised from the quadrant opposite to the bleached area. 50nm thick sections were obtained using an Ultracut ultramicrotome (UCT, Leica) equipped with a Ultra 45 diamond knife (Ultra 45, DiATOME, Hatfield, USA) and collected on formvar-coated 200-mesh copper grids (EMS). Sections were counterstained with uranyl acetate and lead citrate and imaged in a FEI Tecnai G20 Transmission Electron Microscope (FEI, Hillsboro, USA) operated at 80-200 keV and equipped with a Veleta 2K x 2K CCD camera (Olympus, Hamburg, Germany). Around 200 electron micrographs were collected (pixel size = 0.7nm) for each sample. Data were analyzed blindly using the ImageJ software. Double-membraned structures per presynaptic terminal were counted.

### Electrophysiology

Whole cell patch-clamp recordings were performed on autaptic hippocampal neurons at 13–18 DIV. All recordings were obtained at ~25°C from neurons clamped at −70 mV with a Multiclamp 700B amplifier (Molecular Devices, Sunnyvale, USA) under the control of Clampex 10.4 software (Molecular Devices). Data were sampled at 10kHz and low-pass Bessel filtered at 3kHz. Series resistance was compensated at 70% and cells whose series resistance changed more than 25% throughout the recording session were excluded from the analysis. Neurons were immersed in standard extracellular solution consisting of 140mM NaCl, 2.4mM KCl, 10mM HEPES, 10mM glucose, 2mM CaCl_2_ and 4mM MgCl_2_. The borosilicate glass electrodes (3-8 MΩ) were filled with the internal solution containing 136mM KCl, 17.8mM HEPES, 1mM EGTA, 0.6mM MgCl_2_, 4mM ATP-Mg, 0.3mM GTP-Na, 12mM phosphocreatine, and 50U/ml phosphocreatine kinase. All solutions were adjusted to pH 7.4 and osmolarity of ~300mOsm.

Coverslips with cultured neurons were placed on Olympus IX73 microscope (Olympus, Hamburg, Germany) with 20x phase contrast objective. For Supernova bleaching, illumination from a Mercury Vapor Short Arc lamp (X-Cite 120PC Q, Excelitas Technologies, Waltham, USA) was filtered through a 560/40nm filter cube (Olympus U-FF mCherry) and controlled with a mechanical shutter. Lamp iris settings (100%) resulted in 71% bleaching of Supernova intensity (as compared to 22% of GFP bleaching), after 60 seconds of illumination.

From each neuron, 6 sweeps of EPSCs were evoked with a 2ms voltage step from -70mV to 0mV at 0.2Hz. 60 seconds illumination started immediately after end of 6^th^ sweep, and after 5 min of waiting second EPSC was recorded. Control condition without illumination included 6 min waiting period. During recordings with wortmannin, 1μM wortmannnin solution was applied onto the cell using a fast-flow system from the beginning of first EPSC until end of recording session. Electrophysiological data were analyzed offline using Axograph X (Axograph Scientific, Berkeley, USA), Excel (Microsoft, Redmond, USA) and Prism (GraphPad, La Jolla, USA).

### Image acquisition and quantification

All images were acquired on a spinning disc confocal microscope (Zeiss Axio Oberserver.Z1 with Andor spinning disc and cobolt, omricron, i-beam laser) (Zeiss, Oberkochen, Germany) using either a 40x or 63x 1.4 NA Plan-Apochromat oil objective and an iXon ultra (Andor, Belfast, UK) camera controlled by iQ software (Andor, Belfast, UK).

Images were processed using ImageJ and OpenView software (written by Dr. Noam Ziv, Technion Institute, Haifa, Israel). In brief with the OpenView software, multi-channel intensities were measured using a box routine associated with individual boutons. Boxes varied between 7×7 and 9×9 pixel in size, whereas settings were kept the same (e.g. thresholds). The average intensity (synaptic proteins, eGFP-LC3 et cetera) was calculated from all picked puncta and normalized to the control (untreated or unbleached).

For quantification of # of puncta separated axons were randomly picked and the number of puncta per unit length was counted manually. For Supernova experiments, axons were selected from live images showing no or little eGFP-LC3 staining. All Supernova evaluations were normalized to the unbleached control.

To determine the fraction of extrasynaptic eGFP-LC3 puncta positive for Syp-SN/Syn-SN/Syt-SN/Synaptophysin1/Synapsin1/Synaptotagmin1, multi-channel images were manually scanned for eGFP-LC3 puncta within the bleached area that were not colocalizing with Bassoon. Out of these extrasynaptic eGFP-LC3 puncta, the fraction of eGFP-LC3 puncta positive for a specific synaptic protein was quantified.

For FM dye uptake intensities, images of the Syp/Syn-Supernova signal taken before bleaching were used as a mask to define Syp/Syn-Supernova positive puncta. Afterwards, FM 1-43 intensity in Syp/Syn-Supernova positive puncta was quantified using OpenView.

### Experimental Design and Statistical Analyses

Statistical design for all experiments can be found in the figure legends. Independent experiments equal independent cultures. All data representations and statistical analyses were performed with Graph-pad Prism.

## Results

### Monitoring presynaptic autophagy

The primary goal of this study is to examine how and whether the local generation of ROS around SVs triggers a synaptic clearance response that removes damaged proteins. Based on previous studies showing that elevated ROS levels around organelles such as mitochondria leads to the activation of autophagy (Ashrafi et al., 2014; Wang et al., 2012; Yang and Yang, 2011), we anticipated that a similar generation of ROS around SVs may also induce a presynaptic autophagy based clearance program. Thus, while other clearance mechanisms, such as the endo-lysosomal or the proteasome system, could also be activated (see below), we initially sought to develop a live-cell imaging based system that could detect changes in presynaptic autophagy, following different insults.

To achieve this goal, we initially created a lentiviral vector (FU-Syp-mCh-P2A-eGFP-LC3) that co-expresses mCherry-tagged Synaptophysin (Syp-mCh), as a presynaptic marker, and eGFP-tagged LC3, to detect autophagic vacuoles (AVs) (Figure 1A). To allow the independent expression of Syp-mCh and eGFP-LC3, a P2A cleavage site was placed between the two coding sequences (Figure 1A). The vector was then tested in a number of different assays. First, it was lentivirally transduced into Hela cells, where Syp-mCh and eGFP-LC3 both exhibited a largely diffuse cytoplasmic distribution (Figure 1B). The addition of 100μM chloroquine that impedes autophagic flux by blocking the fusion of lysosomes with autophagosomes (Galluzzi et al., 2016; Klionsky et al., 2012), resulted in a redistribution of eGFP-LC3 into a punctate pattern that colocalizes with endogenous LC3 and the autophagophore marker p62 (Johansen, 2011) (Figure 1B). However, Syp-mCh retained its diffuse cytosolic pattern and was not recruited into AVs (Figure 1B). These data indicate not only that the P2A site is efficiently cleaved, but also that the eGFP-LC3 portion of the vector reliably reports the formation of AVs as previously reported (Klionsky et al., 2012; Mizushima et al., 2010; Okerlund et al., 2017).

**Figure 1.**
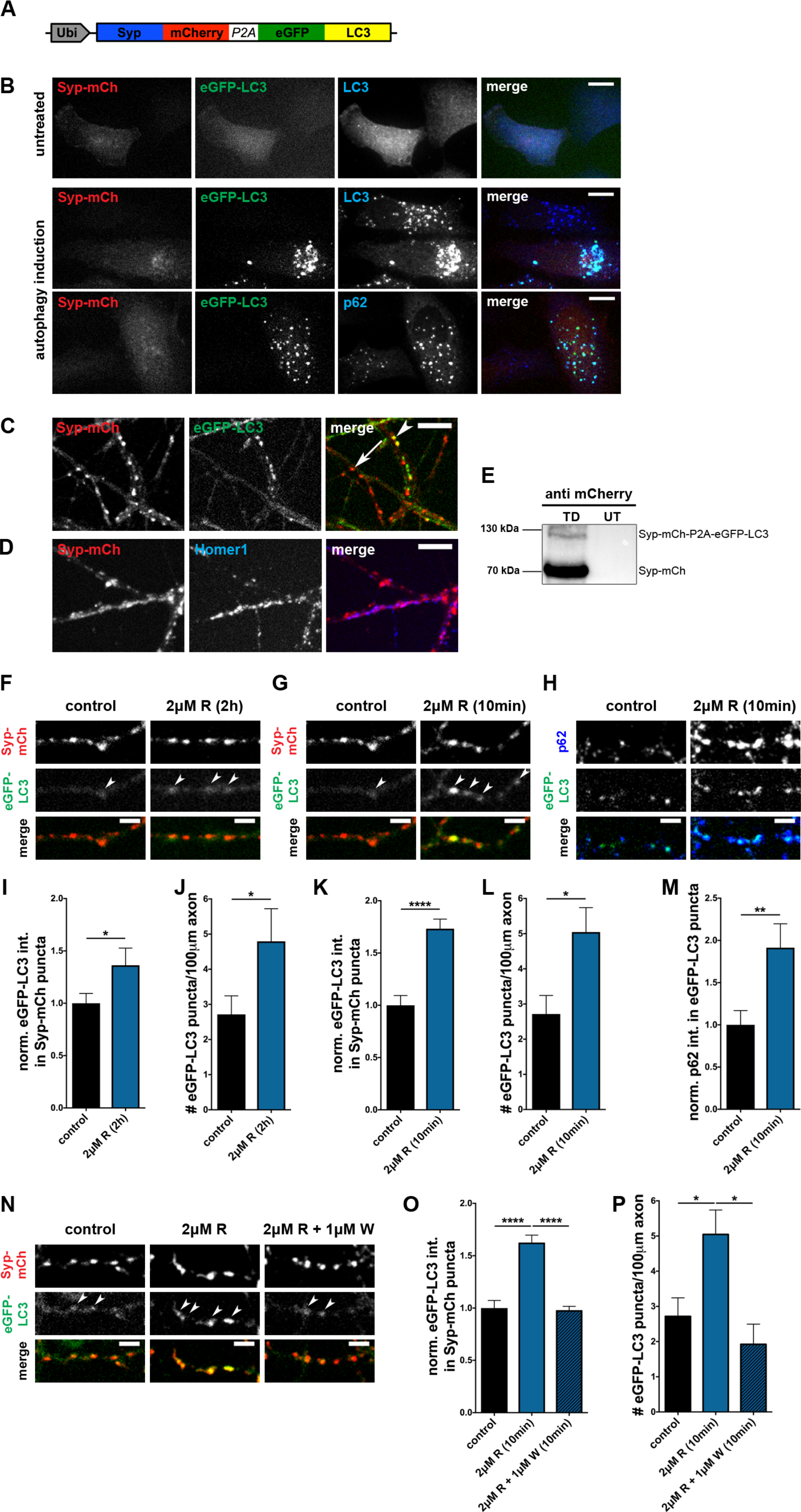
Rapamycin induces rapid increase in presynaptic autophagy. (A) Schematic of lentiviral vector FU-Syp-mCherry-P2A-eGFP-LC3 expressing Synaptophysin (Syp)-mCherry and eGFP-LC3 under an ubiquitin promoter. P2A cleavage site separates the two proteins. (B) Autophagy induction (EBSS + 100μM chloroquine for 2 hours) of FU-Syp-mCherry-P2A-eGFP-LC3 expressing HeLa cells, demonstrating that following autophagy induction eGFP-LC3 puncta colocalize with both endogenous LC3 and p62, but not Syp-mCherry. (C) Live-cell images of cultured hippocampal neurons expressing with FU-Syp-mCherry-P2A-eGFP-LC3 at 2 DIV and analyzed at 14 DIV. Syp-mCherry and eGFP-LC3 exhibit different patterns indicating P2A mediated cleavage (arrow indicates Syp-mCherry puncta, arrowhead indicates colocalization of Syp-mCherry and eGFP-LC3). (D) Representative images of hippocampal neurons infected with FU-Syp-mCherry-P2A-eGFP-LC3 and immunostained with antibodies against the postsynaptic protein Homer1. Colocalization of Syp-mCherry and Homer1 indicate presynaptic targeting of Syp-mCherry. (E) Western Blot of lysates from hippocampal neurons infected (TD) or un-infected (UT) with FU-Syp-mCherry-P2A-eGFP-LC3 and stained with mCherry antibodies. Upper band: uncleaved Syp-mCherry-eGFP-LC3 fusion protein. Lower band: cleaved Syp-mCherry. High ratio of Syp-mCh/Syp-mCh-P2A-eGFP-LC3 band indicate efficient cleavage. (F-H) Images of hippocampal neurons expressing FU-Syp-mCherry-P2A-eGFP-LC3, treated with 2μM rapamycin for 2 hours (F) or 10 min (G and H) before fixation (F-H) and staining with antibodies against p62 (H). (I-M) Quantification of the normalized intensity of eGFP-LC3 levels at Syp-mCherry puncta (I and K) as well as the number of puncta/100 μM of axon (J and L) after 2 hours (I and J) or 10 min (K and L) of 2μM rapamycin treatment. (I: control = 1 ± 0.094, n = 412 synapses, 3 independent experiments; 2μM R (2h) = 1.36 ± 0.164, n = 301 synapses, 3 independent experiments, p=0.0414). (J: control = 2.72 ± 0.529, n = 40 axons, 4 independent experiments; 2μM R (2h) = 4.80 ± 0.928, n = 20 axons, 2 independent experiments, p=/0.0407). (K: control = 1 ± 0.094, n = 412 synapses, 3 independent experiments; 2μM R (10min) = 1.73 ± 0.092, n = 343 synapses, 3 independent experiments; p<0.0001). (L: control = 2.72 ± 0.529, n = 40 axons, 4 independent experiments; 2μM R (10min) = 5.05 ± 0.695, n = 47 axons, 4 independent experiments; p=0.0111). Quantification of the normalized p62 levels at eGFP-LC3 puncta (M) (M: control = 1 ± 0.170, n = 50 puncta, 3 independent experiments; 2μM R (10min) = 1.91 ± 0.283, n = 52 puncta, 3 independent experiments; p=0.0072) confirming that eGFP-LC3 puncta depict autophagic organelles. (N) Images of hippocampal neurons infected with FU-Syp-mCherry-P2A-eGFP-LC3 and treated with 1μM wortmannin prior and during a 10 min incubation with 2μM rapamycin. (O and P) Quantification of (N) showing that wortmannin suppresses the induction of autophagy at Syp-mCherry puncta (O) and along axons (P) following the addition of rapamycin (O: control = 1 ± 0.073, n = 540 synapses, 4 independent experiments; 2μM R (10min) = 1.63 ± 0.071, n = 469 synapses, 4 independent experiments; 2μM R + 1μM W (10min) = 0.98 ± 0.036, n = 152 synapses, 2 independent experiments, p<0.0001 and p<0.0001). (P: control = 2.72 ± 0.529, n = 40 axons, 4 independent experiments; 2μM R (10min) = 5.05 ± 0.695, n = 47 axons, 4 independent experiments; 2μM R + 1μM W (10min) = 1.92 ± 0.573, n = 20 axons, 2 independent experiments, p=0.0187 and p=0.01). Scale bars: 10μm (B, C and D) and 5μm (F, G, H and N). Error bars represent SEM. Unpaired T-test (I, J, K, L and M) and ANOVA Tukey’s multiple comparisons test (O and P) was used to evaluate statistical significance.

In a second set of experiments, we examined whether Syp-mCh faithfully labeled presynaptic sites. Here, dissociated cultures of hippocampal neurons were infected to 30% with our lentiviral vector (FU-Syp-mCherry-P2A-eGFP-LC3) at 2-3 days in vitro (DIV) and analyzed by immuno-cytochemistry at 13-15 DIV. Immuno-staining fixed cultures with antibodies to the postsynaptic density (PSD) protein Homer1 revealed that Syp-mCh forms puncta along the cell somas and dendrites of un-infected cells that colocalize with Homer1 puncta (Figure 1D), consistent with the presynaptic localization of other XFP-tagged Synaptophysin as reported previously (Li et al., 2010). A comparison of Syp-mCh and eGFP-LC3 signals in primary hippocampal neurons during live cell imaging reveals that a small fraction (~10%) of the Syp-mCh positive puncta colocalizes with eGFP-LC3 positive puncta (Figure 1C). This minimal colocalization suggests that the P2A site is functioning properly to uncouple these two proteins. This concept is further supported by western blots of cellular lysates of infected hippocampal neurons stained with a mCherry antibody. Here, greater than 95% of the immuno-reactivity is present in the 70kDa Syp-mCh band versus the uncleaved 120kDa Syp-mCh-P2A-eGFP-LC3 band (Figure 1E), supporting the conclusion that once expressed in neurons each reporter is free to operate independently.

In a third set of experiments, we examined how the induction of autophagy with 2μM rapamycin (Boland et al., 2008; Hernandez et al., 2012; Spilman et al., 2010) affected the distribution of eGFP-LC3 relative to Syp-mCh in neurons. Initially, rapamycin was added to sparsely FU-Syp-mCherry-P2A-eGFP-LC3 infected hippocampal cultures (13-15 DIV) for 2 hours, as most previously studies had shown that this condition can induce autophagy in neurons (Hernandez et al., 2012). To identify ‘synaptic’ changes in eGFP-LC3 levels, we analyzed the average intensities of eGFP-LC3 puncta that colocalized with Syp-mCh puncta in fixed neurons. This revealed a modest (36%) but significant increase in eGFP-LC3 intensities within presynaptic boutons compared to untreated control neurons (Figure 1F and I). Monitoring the number of eGFP-LC3 puncta per unit length of axon revealed that 2 hours of rapamycin treatment significantly increased the number of eGFP-LC3 puncta present in axons compared to non-treated control neurons (Figure 1F and J). These data are consistent with the concept that rapamycin can induce the formation of autophagosomes/AVs in hippocampal axons. However, given that vesicular transport is quite rapid, it is unclear whether during the 2 hour period the newly formed AVs arose at synapses and dispersed into the axons and/or were generated within axons and then accumulate within presynaptic boutons. We thus explored whether AVs would appear in as little as 10 minutes following the addition of rapamycin. Surprisingly, we found that not only did eGFP-LC3 puncta appear in axons during this short period of induction (Figure 1G and L), but eGFP-LC3 intensity was dramatically increased within presynaptic boutons marked with Syp-mCh (Figure 1G and K). Importantly, we also found that appearing eGFP-LC3 puncta were positive for the autophagy cargo receptor p62 (Johansen, 2011) (Figure 1H and M), suggesting that they are indeed autophagosomes and are forming locally within presynaptic boutons. To further explore whether the observed rapamycin induced AV formation at synapses is induced via the conventional autophagy pathway, which includes the PI3K Vps34 (Lilienbaum, 2013; Rubinsztein et al., 2012), we included 1μM wortmannin, a PI3K inhibitor (Carpenter and Cantley, 1996; Klionsky et al., 2012), together with rapamycin during the 10 minutes incubation period. This manipulation abolished the accumulation of eGFP-LC3 puncta in both presynaptic boutons (marked with Syp-mCh) (Figure 1N and O) and along axons (Figure 1N and P). Taken together these data indicate that the machinery necessary for the rapid generation of AVs is located within or very near to presynaptic boutons and can be triggered by a PI3K-dependent pathway.

### Light-induced ROS generation triggers presynaptic autophagy

The ability of rapamycin to induce presynaptic autophagy within 10 minutes strongly suggests that presynaptic boutons contain local clearance mechanisms, such as autophagy, that could in principle deal with locally damaged proteins in real-time. As a direct test of this hypothesis, we explored whether the real-time damage of SV proteins via the production of reactive oxygen species (ROS) (Takemoto et al., 2013) could also trigger the rapid clearance of these molecules via, e.g. autophagy. To accomplish this goal, we made use of a molecular variant of GFP called Supernova, a monomeric version of KillerRed (Bulina et al., 2006), previously shown to generate ROS following its excitation with 550-590nm light (Takemoto et al., 2013). As other photosensitizers, short-lived ROS generated by Supernova is expected to damage proteins within 1-4nm of the source (Linden et al., 1992; Takemoto et al., 2013). Thus to restrict the actions of the ROS to SVs, we initially fused Supernova to the short cytoplasmic tail of the SV protein Synaptophysin (creating Synaptophysin-Supernova; Syp-SN). This was then subcloned and co-expressed with eGFP-LC3 via our lentiviral vector (FU-Syp-Supernova-P2A-eGFP-LC3) (Figure 2A) (see also Figure 6A).

**Figure 2.**
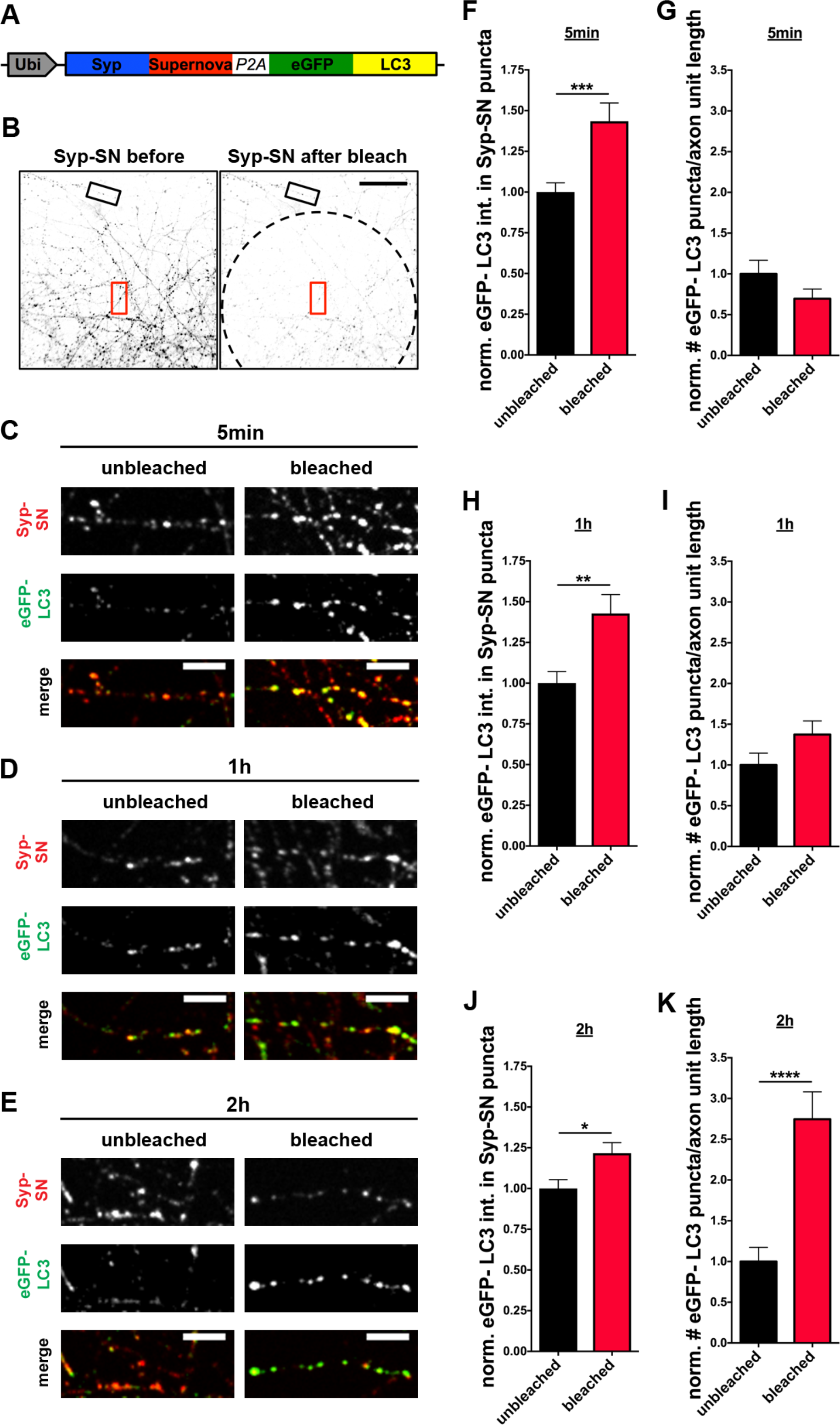
Rapid induction of autophagy by ROS mediated damage by Synaptophysin-Supernova. (A) Schematic of FU-Syp-Supernova-P2A-eGFP-LC3 expression vector. (B) Low-magnification images of hippocampal neurons expressing FU-Syp-Supernova-P2A-eGFP-LC3 grown on top of uninfected neurons before and after photobleaching a region of interest (dashed line). Boxes represent areas within (red) and outside (black) bleached area used for analysis. (C-E) Images of axon segments (5 min, 1 hour and 2 hours after bleaching) that were subsequently fixed and stained with antibodies against GFP to detect eGFP-LC3 and Supernova to detect Syp-SN. Data indicate that autophagy at synapses can be rapidly induced through Syp-SN photobleaching. (F) Quantification of normalized eGFP-LC3 intensities within Syp-SN puncta 5 min (C) after bleaching (unbleached = 1 ± 0.057, n = 119 synapses, 3 independent experiments; bleached = 1.43 ± 0.113, n = 132 synapses, 3 independent experiments, p=0.001). (G) Quantification of the normalized number of eGFP-LC3 puncta per unit axon length, in axons 5 min after photobleaching (C) (unbleached = 1 ± 0.166, n = 17 axons, 3 independent experiments; bleached = 0.70 ± 0.119, n = 18 axons, 3 independent experiments). (H) Quantification of normalized eGFP-LC3 intensities within Syp-SN puncta 1 hour (D) after bleaching (unbleached = 1 ± 0.071, n = 132 synapses, 3 independent experiments; bleached = 1.43 ± 0.117, n = 167 synapses, 3 independent experiments, p=0.0035). (I) Quantification of the normalized number of eGFP-LC3 puncta per unit axon length, in axons 1 hour after photobleaching (D) (unbleached = 1 ± 0.146, n = 24 axons, 3 independent experiments; bleached = 1.37 ± 0.166, n = 24 axons, 3 independent experiments). (J) Quantification of normalized eGFP-LC3 intensities within Syp-SN puncta 2 hours (E) after bleaching (unbleached = 1 ± 0.054, n = 136 synapses, 3 independent experiments; bleached = 1.22 ± 0.065, n = 141 synapses, 3 independent experiments, p=0.0111). (K) Quantification of the normalized number of eGFP-LC3 puncta per unit axon length, in axons 2 hours after photobleaching (E) (unbleached = 1 ± 0.173, n = 23 axons, 3 independent experiments; bleached = 2.75 ± 0.336, n = 22 axons, 3 independent experiments, p<0.0001). Scale bars: 50μm (B), 10μm (C, D and E). Error bars represent SEM. Unpaired T-test was used to evaluate statistical significance.

As with the FU-Syp-mCherry-P2A-eGFP-LC3 vector, we then verified that both the Syp-SN and eGFP-LC3 portions of the vector were expressed and processed. We also verified that the eGFP-LC3 segment was recruited to p62 positive AVs in HeLa cells treated with chloroquine and that Syp-SN properly localized at Homer1 positive synapses as Syp-mCh (data not shown). Moreover, we confirmed in Hela cells that 80% of the Supernova fluorescence could be photobleached during a 60 seconds exposure of 581nm wavelength light from a mercury lamp, and verified that ROS was being generated with the superoxide indicator Dihydroethidium (DHE), as previously shown (Takemoto et al., 2013).

To explore whether a local increase in ROS production near SVs can induce presynaptic autophagy, primary hippocampal neurons grown on μ-Slide 8 Well culture dishes were sparsely infected with FU-Syp-Supernova-P2A-eGFP-LC3 at 2-3 DIV. Around 14 DIV, they were transferred to a spinning disc confocal microscope equipped with a temperature controlled live-cell imaging chamber. Prior to bleaching selected fields of view, axons from infected neurons growing on top of uninfected neurons were selected and imaged during excitation with a 491nm (for the eGFP-LC3 signal) and a 561nm laser (for the Syp-SN signal). Subsequently, a subregion, selected with a field diaphragm, was bleached by exposing cells to 581nm light from a mercury lamp for 60 seconds (Figure 2B), a condition found to bleach approximately 80% of the initial fluorescence. Cultures were fixed 5-120 minutes post bleaching and immuno-stained with antibodies against GFP and Supernova, allowing the post-hoc identification of synapses within and outside of the bleached area and the levels and redistribution of eGFP-LC3. Comparing the intensity of eGFP-LC3 at Syp-SN positive puncta within and outside the bleached area revealed a significant increase in synaptic eGFP-LC3 intensity in the bleached area within 5 minutes of initial bleaching (Figure 2C and F). However, at that time point, the number of eGFP-LC3 puncta per axon unit length is not changed compared to the unbleached control (Figure 2C and G). Similarly, 1 hour after bleaching, eGFP-LC3 levels are still elevated in Syp-SN positive synapses inside the bleached area compared to outside, with only a modest increase in the number of eGFP-LC3 puncta per unit length of axon (Figure 2D, H and I). Intriguingly, 2 hours after triggering ROS production, eGFP-LC3 levels remain somewhat elevated at Syp-SN positive synapses, and dramatically accumulated as small puncta along axons inside the bleached area (Figure 2E, J and K) compared to those outside. These latter data imply that the synaptic increase in ROS rapidly induces presynaptic autophagy and that subsequent flux carries the autophagosomal membranes into axons.

To investigate whether the observed ROS-induced increase in eGFP-LC3 puncta is dependent on the conventional PI3K/Vps34 autophagy pathway, 1μM wortmannin was added to neurons before bleaching SN and maintained in the culture for the following 2 hours, after which neurons were fixed and analyzed. In cells that were not treated with wortmannin, eGFP-LC3 intensity within presynaptic boutons as well as the number of eGFP-LC3 puncta per unit length of axon remained elevated (Figure 4A, C and D) compared to the unbleached control. In contrast, the inclusion of wortmannin was found to inhibit the light-induced increase in eGFP-LC3 intensity within presynaptic boutons (Figure 3B and E), but had no effect on the number of eGFP-LC3 puncta per unit length of axon (Figure 3B and F). These data indicate that the ROS-induced increase in presynaptic autophagy may be dependent on the PI3K signaling pathway, while autophagy within axons is not. Note that while most of the Syp-SN is synaptic (data not shown), extrasynaptic pools are likely present, presumably engaged in the active transport within mobile pools of SVs (Cohen et al., 2013; Maas et al., 2012; Tsuriel et al., 2006). Photobleached damage of this pool could thus contribute to a PI3K-independent form of axonal autophagy in axons, as already described for other cell types (Chu et al., 2007; Lemasters, 2014; Zhu et al., 2007).

**Figure 3.**
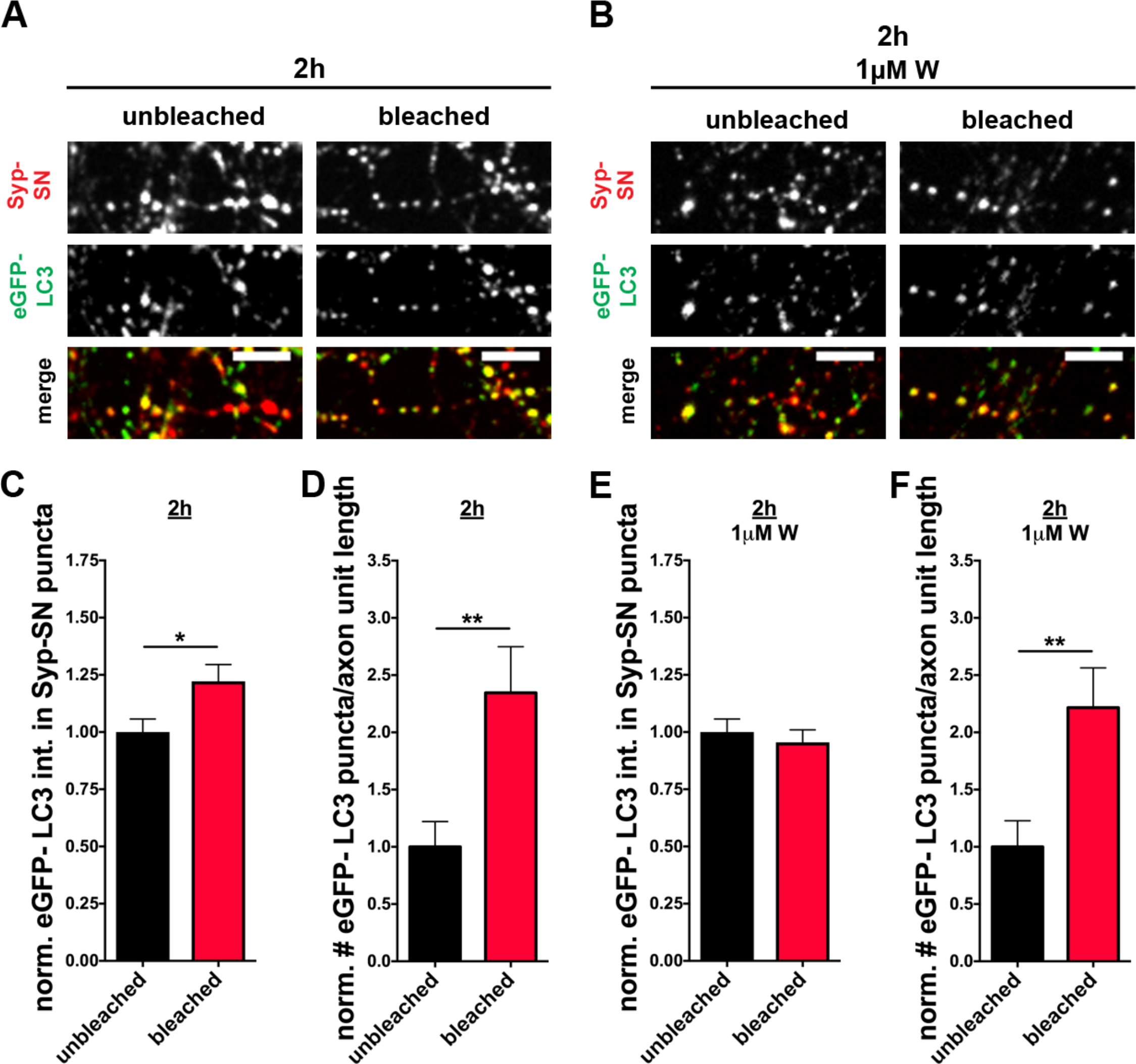
ROS induced increase in eGFP-LC3 levels at presynaptic boutons is PI3K-dependent. (A and B) Images of hippocampal axons/synapses expressing FU-Syp-Supernova-P2A-eGFP-LC3 that were fixed and stained with antibodies against GFP and Supernova 2 hours after photobleaching, either in the absence (A) or presence of 1μM wortmannin (B). (C and D) Quantification of normalized eGFP-LC3 intensities within Syp-SN puncta (C) or the normalized number of eGFP-LC3 puncta per unit axon length (D) in culture not treated with wortmannin. (C: unbleached = 1 ± 0.057, n = 174 synapses, 3 independent experiments; bleached = 1.22 ± 0.073, n = 174 synapses, 3 independent experiments, p=0.0173) (D: unbleached = 1 ± 0.221, n = 19 axons, 3 independent experiments; bleached = 2.35 ± 0.403, n = 21 axons, 3 independent experiments, p=0.0071). (E and F) Quantification of normalized eGFP-LC3 intensities within Syp-SN puncta (E) or number of eGFP-LC3 puncta per unit axon length (F) in culture treated with wortmannin before and after photobleaching. (E: unbleached = 1 ± 0.057, n = 179 synapses, 3 independent experiments; bleached = 0.95 ± 0.055, n = 164 synapses, 3 independent experiments). (F: unbleached = 1 ± 0.228, n = 18 axons, 3 independent experiments; bleached = 2.22 ± 0.348, n = 21 axons, 3 independent experiments, p=0.0077). Scale bars: 10μm. Error bars represent SEM. Unpaired T-test was used to evaluate statistical significance.

**Figure 4.**
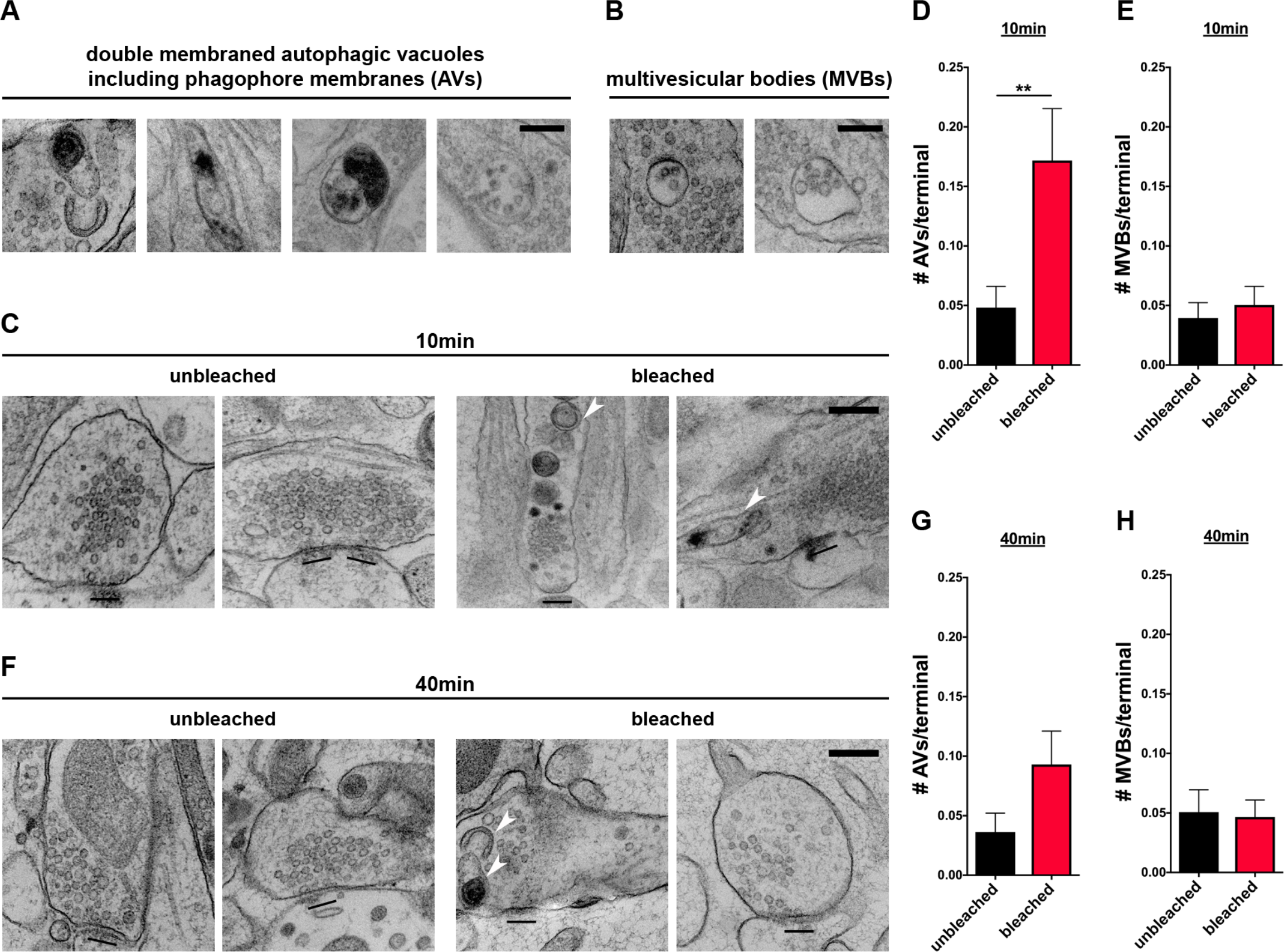
Syp-SN mediated ROS production increases autophagic vacuoles (AVs) in presynaptic terminals. (A and B) Example EM micrographs of organelles quantified as autophagic vacuoles (AVs) (A) or multivesicular bodies (MVBs) (B). (C and F) Representative EM micrographs of bleached or unbleached synapses 10 (C) or 40 (F) min after photobleaching. Arrowheads indicate double membraned AVs. Note, # of AVs but not MVBs is significantly increased 10 min following Syp-SN mediated ROS production. (D and E) Quantification of the number of AVs (D) or MVBs (E) per terminal 10 min after photobleaching (D: unbleached = 0.05 ± 0.018, n = 228 synapses, 1 independent experiments; bleached = 0.17 ± 0.044, n = 198 synapses, 2 independent experiments, p=0.0062) (E: unbleached = 0.04 ± 0.013, n = 228 synapses, 2 independent experiments; bleached = 0.05 ± 0.016, n = 198 synapses, 2 independent experiments). (G and H) Quantification of the number of AVs (G) or MVBs (H) per terminal 40 min after photobleaching (G: unbleached = 0.04 ± 0.016, n = 138 synapses, 2 independent experiments; bleached = 0.09 ± 0.028, n = 215 synapses, 2 independent experiments) (H: unbleached = 0.05 ± 0.019, n = 138 synapses, 2 independent experiments; bleached = 0.05 ± 0.014, n = 215 synapses, 2 independent experiments). Scale bars: 300nm (C and F), 200nm (A and B). Error bars represent SEM. Unpaired T-test was used to evaluate statistical significance.

### ROS induced damage to Synaptophysin promotes AV formation

The appearance of eGFP-LC3 positive puncta within the axons and presynaptic boutons of Synaptophysin-Supernova expressing cells following photobleaching suggests that this insult induces the autophagic clearance of damaged SVs and their proteins. To formally test this hypothesis, we performed transmission electron microscopy of FU-Syp-Supernova-P2A-eGFP-LC3 infected hippocampal neurons. Infected neurons grown on sapphire disks were photobleached with 581nm light from a mercury lamp for 60 seconds. Similar to our live imaging experiments, a field diaphragm was used to create bleached and unbleached regions on the same sapphire disk before high pressure freezing and further processing for EM. The number of double-membraned organelles (autophagic vacuoles = AVs) within presynaptic boutons or SVs containing axonal varicosities were then quantified as performed previously (Okerlund et al., 2017). Consistent with light level studies (Figure 2F), significantly more AVs per presynaptic terminal were observed 10 minutes after light-induced Synaptophysin damage within the bleached area compared to the unbleached area (Figure 4C and D). Images analyzed ~40 minutes after bleaching revealed a slight but non-significant increase in AVs/terminal (Figure 4F and G). These data indicate that most newly formed autophagosomes quickly leave the synapse.

Conceptually, local ROS induced damage of synaptic proteins could induce not only autophagy but also other degradative pathways such as the endo-lysosomal system. One hallmark of the endo-lysosomal system is the appearance of multivesicular bodies (MVBs) (Ceccarelli et al., 1973; Raiborg and Stenmark, 2009). We thus examined whether the light-induced damage of Synaptophysin also induces the endo-lysosomal pathway by quantifying the presence of synaptic MVBs within photobleached presynaptic boutons by EM. Intriguingly, no change in their number was observed either 10 or 40 minutes after photobleaching compared to unbleached boutons (Figure 4E and H), indicating that the ROS mediated damage of Synaptophysin primarily triggers the activation of autophagy. To confirm this observation, we also monitored whether markers of the endo-lysosomal pathway accumulated in presynaptic boutons following light-induced damage of Synaptophysin. Strikingly, level of the late endosome marker Rab7 are increased at presynaptic boutons 5 minutes after bleaching (Figure 5B and E), and stay elevated compared to the unbleached control for at least 2 more hours (Figure 5E, G and I). Since Rab7 is also abundant on autophagosomes, we stained for another, more specific, MVB marker Chmp2b, which is part of the ESCRT-III complex (Vingtdeux et al., 2012). Interestingly, Chmp2b also accumulates at boutons 1 hour after bleaching (Figure 5 C and H). These observations indicate that ROS mediated damage to Synaptophysin/SVs may also engage other degradative pathways such as the endo-lysosomal system.

**Figure 5.**
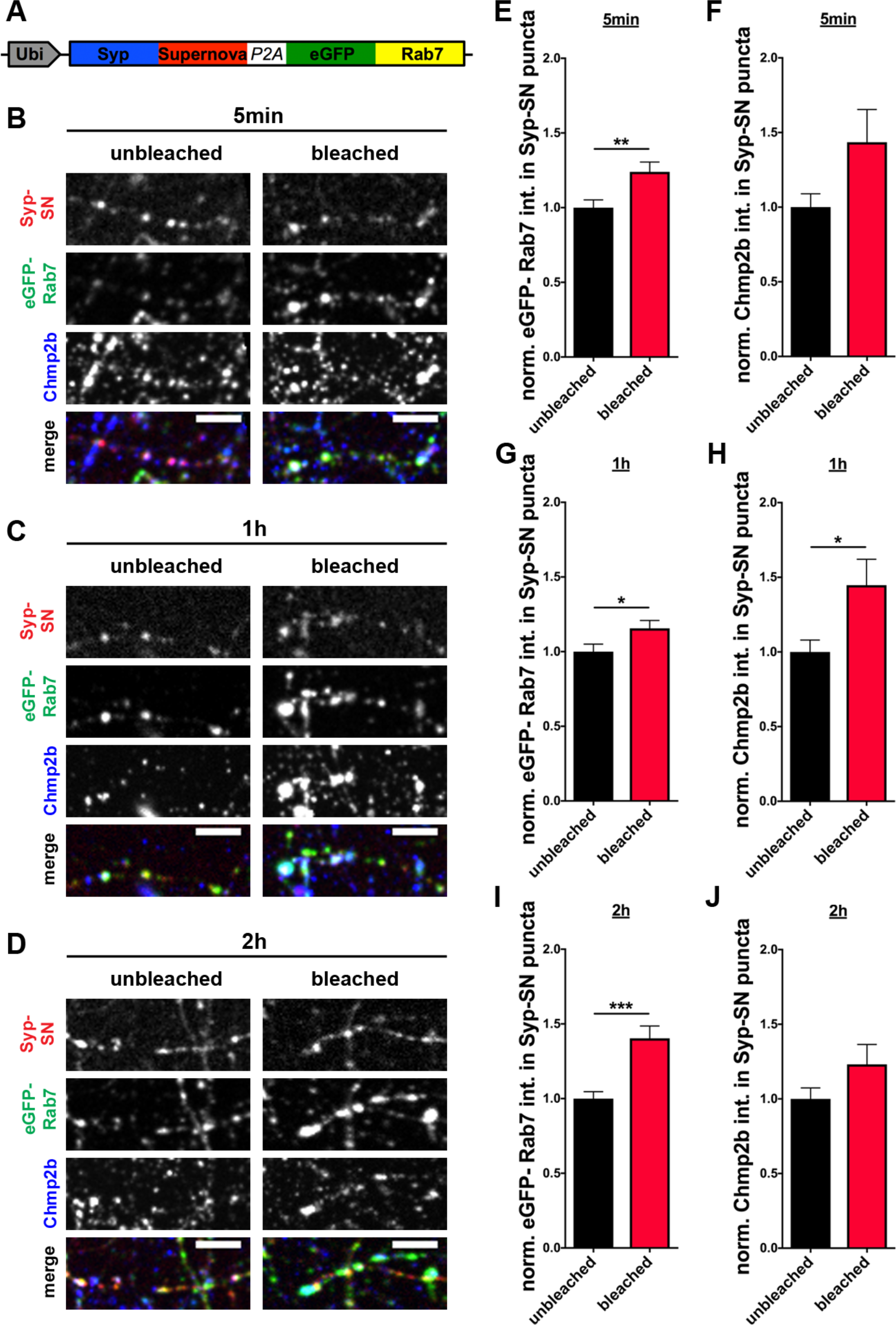
Syp-SN mediated ROS production increases eGFP-Rab7 and Chmp2b levels at presynaptic boutons. (A) Schematic of FU-Syp-Supernova-P2A-eGFP-Rab7 expression vector. (B, C and D) Images of hippocampal axons/synapses expressing FU-Syp-Supernova-P2A-eGFP-Rab7 that were fixed and stained with antibodies against GFP, Supernova and Chmp2b, 5 min, 1 hour and 2 hours after Syp-SN mediated ROS production. (E, G and I) Quantification of the normalized eGFP-Rab7 intensity in Syp-SN puncta 5 min, 1 hour or 2 hours after photobleaching of Syp-SN (E: unbleached = 1 ± 0.052, n = 249 synapses, 4 independent experiments; bleached = 1.24 ± 0.066, n = 314 synapses, 4 independent experiments, p=0.0063) (G: unbleached = 1 ± 0.050, n = 280 synapses, 4 independent experiments; bleached = 1.16 ± 0.053, n = 373 synapses, 4 independent experiments, p=0.0376) (I: unbleached = 1 ± 0.046, n = 258 synapses, 4 independent experiments; bleached = 1.40 ± 0.083, n = 352 synapses, 4 independent experiments, p=0.0001). Note, Rab7 levels are significantly increased at all three times. (F, H and J) Quantification of the normalized Chmp2b intensity in Syp-SN puncta 5 min, 1 hour or 2 hours after photobleaching of Syp-SN. Levels are significantly increased at 1 hour but not 5 min or 2 hours after ROS mediated damage. (F: unbleached = 1 ± 0.089, n = 67 synapses, 2 independent experiments; bleached = 1.44 ± 0.219, n = 71 synapses, 2 independent experiments) (H: unbleached = 1 ± 0.080, n = 91 synapses, 2 independent experiments; bleached = 1.45 ± 0.174, n = 89 synapses, 2 independent experiments, p=0.0196) (J: unbleached = 1 ± 0.074, n = 108 synapses, 2 independent experiments; bleached = 1.23 ± 0.134, n = 118 synapses, 2 independent experiments). Scale bars: 10μm. Error bars represent SEM. Unpaired T-test was used to evaluate statistical significance.

### Induction of presynaptic autophagy requires the association of Supernova with SVs

As ROS generated by illuminating Supernova is anticipated to damage proteins only within 1-4nm of the sources (Jacobson et al., 2008; Takemoto et al., 2013), it seems reasonable to predict that the induction of presynaptic autophagy is linked to the damage of proteins physically associated with SVs, which are then sorted and gathered into the interior of the newly forming autophagophore membrane. If true, the degree of induction would be related to the physical association of proteins with SVs (Figure 6A).

**Figure 6.**
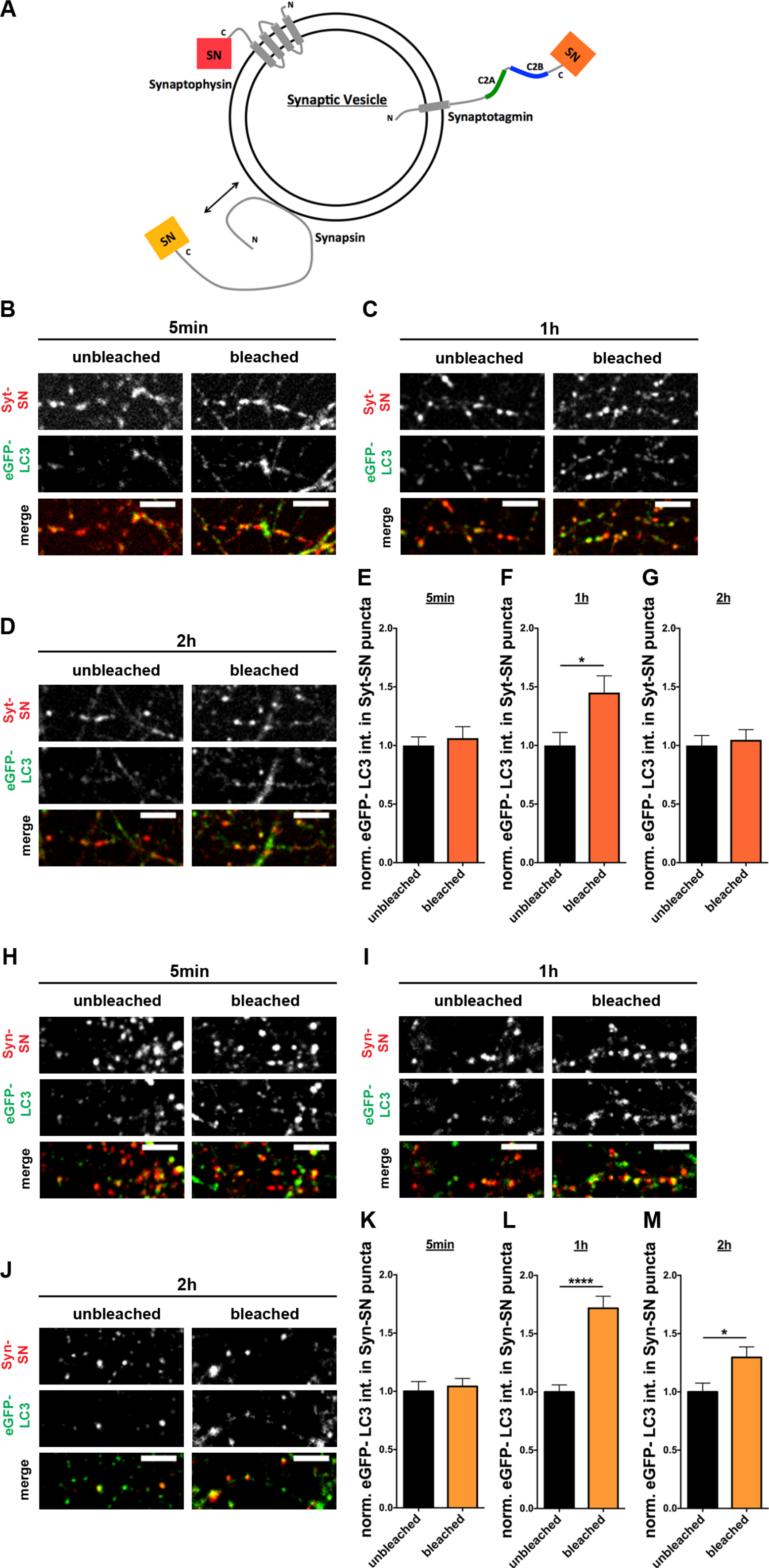
Syt-SN and Syn-SN mediated ROS production increases eGFP-LC3 levels at presynaptic boutons. (A) Schematic of a SV containing Synaptophysin, Synaptotagmin or Synapsin tagged with Supernova. Note, the short (95aa) vs. long (346aa) cytoplasmic tails of Synaptophysin vs. Synaptotagmin, respectively, which could significantly change the distance of Supernova to the SV membrane. Similarly, tagging Supernova to the much larger peripherally associated SV protein Synapsin could also affect its distance to the SV membrane. Moreover, as the association of Synapsin with SVs is dynamically regulated, activity can be used to disassociate it from SVs. (B, C and D) Images of hippocampal synapses expressing FU-Syt-Supernova-P2A-eGFP-LC3 that were fixed and stained with antibodies against GFP and Supernova 5 min (B), 1 hour (C) and 2 hours (D) after bleaching. Note, eGFP-LC3 levels are significantly increased 1 hour after Syt-SN mediated ROS production. (E, F and G) Quantification of normalized eGFP-LC3 intensities within Syt-SN puncta, 5 min (E), 1 hour (F) and 2 hours (G) after bleaching. (E: unbleached = 1 ± 0.074, n = 62 synapses, 2 independent experiments; bleached = 1.06 ± 0.098, n = 76 synapses, 2 independent experiments) (F: unbleached = 1 ± 0.111, n = 73 synapses, 2 independent experiments; bleached = 1.45 ± 0.143, n = 81 synapses, 2 independent experiments, p=0.0156) (G: unbleached = 1 ± 0.085, n = 58 synapses, 2 independent experiments; bleached = 1.05 ± 0.087, n = 68 synapses, 2 independent experiments). (H, I and J) Images of hippocampal synapses expressing FU-Syn-Supernova-P2A-eGFP-LC3 that were fixed and stained with antibodies against GFP and Supernova 5 min (H), 1 hour (I) and 2 hours (J) after bleaching. Note, eGFP-LC3 levels are significantly increased 1 hour and 2 hours after Syn-SN mediated ROS production. (K, L and M) Quantification of normalized eGFP-LC3 intensities within Syp-SN puncta, 5 min (K), 1 hour (L) and 2 hours (M) after bleaching. (K: unbleached = 1 ± 0.084, n = 58 synapses, 3 independent experiments; bleached = 1.04 ± 0.069, n = 81 synapses, 3 independent experiments) (L: unbleached = 1 ± 0.061, n = 77 synapses, 3 independent experiments; bleached = 1.72 ± 0.103, n = 103 synapses, 3 independent experiments, p<0.0001) (M: unbleached = 1 ± 0.075, n = 42 synapses, 3 independent experiments; bleached = 1.30 ± 0.090, n = 71 synapses, 3 independent experiments, p=0.0255). Scale bars: 10μm. Error bars represent SEM. Unpaired T-test was used to evaluate statistical significance.

To test this hypothesis, we coupled Supernova to two additional SV proteins, Synaptotagmin (Syt), an integral membrane protein with a long cytoplasmic tail (Chapman, 2002; Hilfiker et al., 1999), and Synapsin (Syn), a larger cytosolic protein (Figure 6A) that dynamically associates with the outer surface of SVs in an activity dependent manner (Chi et al., 2001; Waites and Garner, 2011), potentially allowing for a more attenuated ROS mediated damage to SVs. To permit the simultaneous detection of presynaptic autophagy, we co-expressed Syt-SN or Syn-SN with eGFP-LC3 via our lentiviral vector (FU-Syt-SN-P2A-eGFP-LC3; FU-Syn-SN-P2A-eGFP-LC3). In control experiments, we confirmed that both Syt-SN and Syn-SN and eGFP-LC3 were appropriately processed and that Syt-SN and Syn-SN retained their ability to become selectively localized to presynaptic boutons (data not shown). As described above for Syp-SN, neurons infected at 2-3 DIV were photobleached at 13-15 DIV for 60 seconds and the intensity of eGFP-LC3 within presynaptic boutons quantified. Interestingly, eGFP-LC3 intensity in Syt-SN and Syn-SN puncta as well as the number of eGFP-LC3 puncta along axons did not change within 5 minutes of photobleaching (Figure 6B, E, H and K) compared to unbleached boutons. However, 1 hour after light-induced damage to either Synaptotagmin or Synapsin, eGFP-LC3 intensity significantly increased within presynaptic boutons immuno-positive for Syt-SN (Figure 6C and F) and Syn-SN (Figure 6I and L). When fixed 2 hours after ROS production, eGFP-LC3 levels remained slightly elevated at bleached Syn-SN positive synapses (Figure 6J and M), but returned to unbleached levels in Syt-SN positive synapse (Figure 6D and G). Taken together, these data indicate that, as Synaptophysin, the local ROS mediated damage to the Synaptotagmin and the SV-associated protein Synapsin can induce presynaptic autophagy, albeit at attenuated slower rates.

Intriguingly previous studies have shown that only about 50% of Synapsin within boutons is physically associated with SVs, while the remainder is soluble (Benfenati et al., 1993; Chi et al., 2001; Leal-Ortiz et al., 2008). Given that autophagy directed cargos are primarily membrane bound, we posited that it is the SV bound form of Synapsin-SN that triggers presynaptic autophagy. To test this hypothesis, we took advantage of the activity dependent regulation of Synapsin to trigger its dissociation from SVs and dispersion out of the synapse, using a high KCl (60mM) stimulation (Chi et al., 2001). Remarkably, when photobleaching was performed during a high KCl stimulus, eGFP-LC3 levels did not increase 1 hour after ROS production at synapses over-expressing Syn-SN compared to unstimulated control (Figure 7B and E). There was also no increase in the number of eGFP-LC3 puncta per axon unit length detectable (Figure 7B and F). These data indicate that the induction of presynaptic autophagy is tightly coupled to ROS damage of SV proteins, and thus associated with the normal clearance of mis-folded or damaged SV proteins.

**Figure 7.**
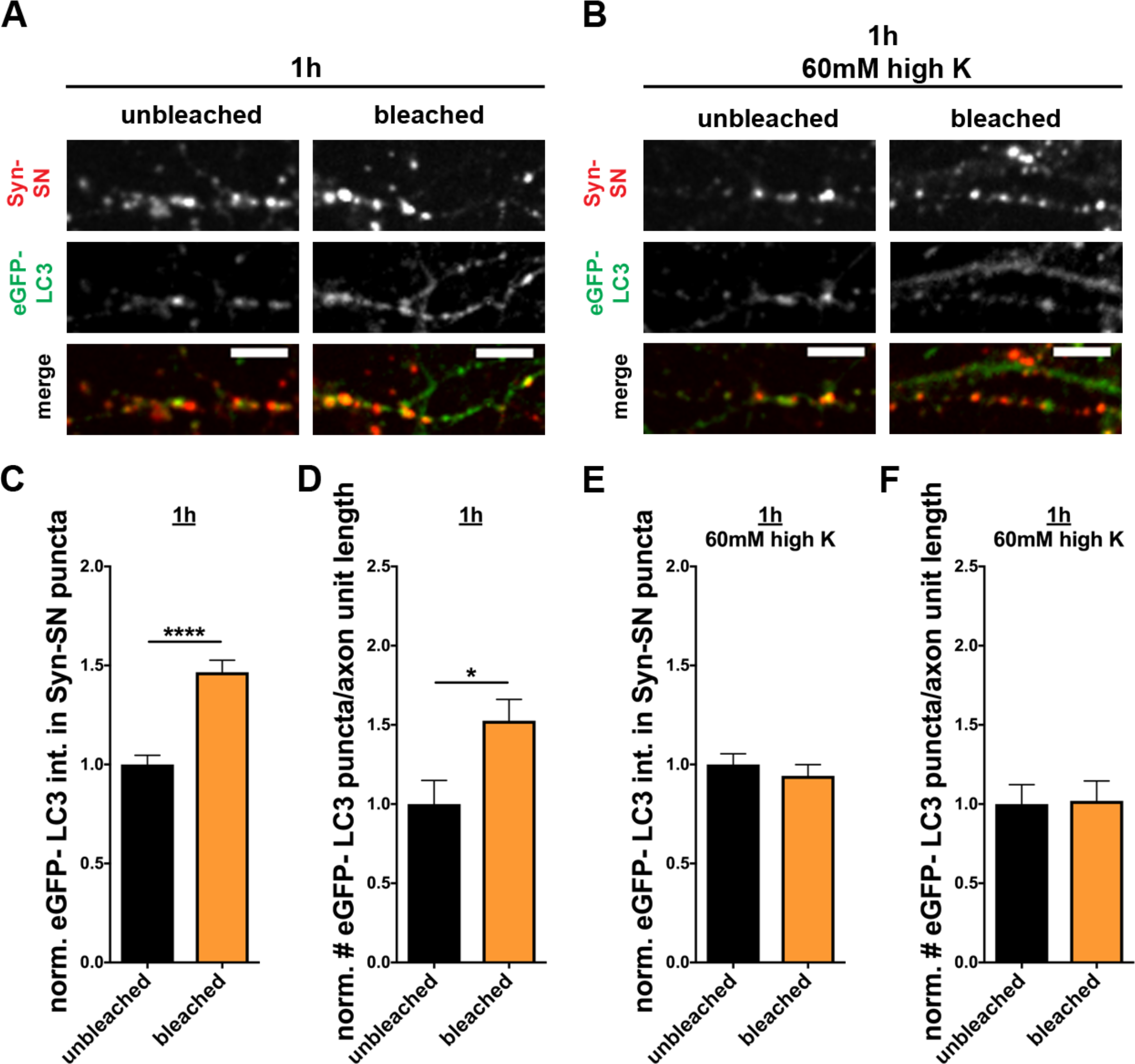
Synapsin dispersion blocks ROS induced increase in eGFP-LC3 levels in axons and boutons expressing Syn-SN. (A and B) Images of hippocampal synapses expressing FU-Syn-Supernova-P2A-eGFP-LC3 that were fixed and stained with antibodies against GFP and Supernova 1 hour after bleaching in the absence (A) or presence of 60mM KCl (B) to increase synaptic activity and induce the dispersion of Syn-SN. (C - F) Quantification of normalized eGFP-LC3 intensities within Syp-SN puncta (C and E) or the normalized number of eGFP-LC3 puncta per unit axon length (D and F) 1 hour after photobleaching of Syn-SN in the absence (C and D) or presence of 60mM KCl (E and F). Note that increased activity to disperse Syn-SN blocks ROS induced increase in axonal and synaptic eGFP-LC3 (C: unbleached = 1 ± 0.047, n = 240 synapses, 3 independent experiments; bleached = 1.47 ± 0.061, n = 330 synapses, 3 independent experiments, p<0.0001) (D: unbleached = 1 ± 0.150, n = 26 axons, 3 independent experiments; bleached = 1.53 ± 0.135, n = 27 axons, 3 independent experiments, p=0.0117) (E: unbleached = 1 ± 0.054, n = 116 synapses, 3 independent experiments; bleached = 0.94 ± 0.057, n = 108 synapses, 3 independent experiments) (F: unbleached = 1 ± 0.123, n = 19 axons, 3 independent experiments; bleached = 1.02 ± 0.126, n = 19 axons, 3 independent experiments). Scale bars: 10μm. Error bars represent SEM. Unpaired T-test was used to evaluate statistical significance.

### Supernova-tagged proteins are more abundant in ROS-induced autophagy organelles than endogenous SV proteins

To date several studies have demonstrated that autophagosomes form in axons upon starvation, rapamycin treatment as well as enhanced synaptic activity (Maday and Holzbaur, 2014; Wang et al., 2015) and become retrogradely transported along the axon towards the soma (Cheng et al., 2015a; Maday et al., 2012). An unresolved question is which synaptic proteins become associated with autophagic cargos. A related question is whether presynaptic autophagy leads to the en-mass removal of SVs or whether can it selectively scavenge damaged proteins. Our ability to damage SV proteins with light and induce autophagy provides a unique opportunity to explore these questions. In an initial experiment, we examined whether Supernova tagged Synaptophysin (Syp-SN) appears in extrasynaptic eGFP-LC3 positive puncta following light-induced ROS production. To distinguish between synaptic and extrasynaptic eGFP-LC3 organelles, cultures were fixed and stained with the presynaptic active zone protein Bassoon and quantified for the fraction of extrasynaptic eGFP-LC3 puncta negative for Bassoon but positive for synaptic proteins 1 hour after bleaching.

In experiments with Syp-SN, we observed that more than 70% of extrasynaptic eGFP-LC3 puncta (also referred to as autophagy cargo organelles) are positive for Syp-SN (Figure 8A and D). This suggests that ROS-damaged Syp-SN is indeed a cargo of these organelles. To investigate whether the presence of Syp-SN in autophagy cargo organelles represents the en-mass engulfment of SVs or the selective removal of this damaged protein, we monitor the distribution of endogenous Synaptotagmin1 within the same neurons, a second core constituent of SVs, in extrasynaptic autophagy organelles following light-induced damage to Synaptophysin-SN. As Synaptotagmin1 is not known to directly interact with Synaptophysin, we reasoned that the ROS mediated damage to Syp-SN would not necessarily damage Synaptotagmin on the same SV. Interestingly, the fraction of extrasynaptic autophagy organelles that are positive for Synaptotagmin1 (Syt1) is dramatically smaller than the fraction of Syp-SN positive autophagy cargo organelles (Figure 8A and D).

**Figure 8.**
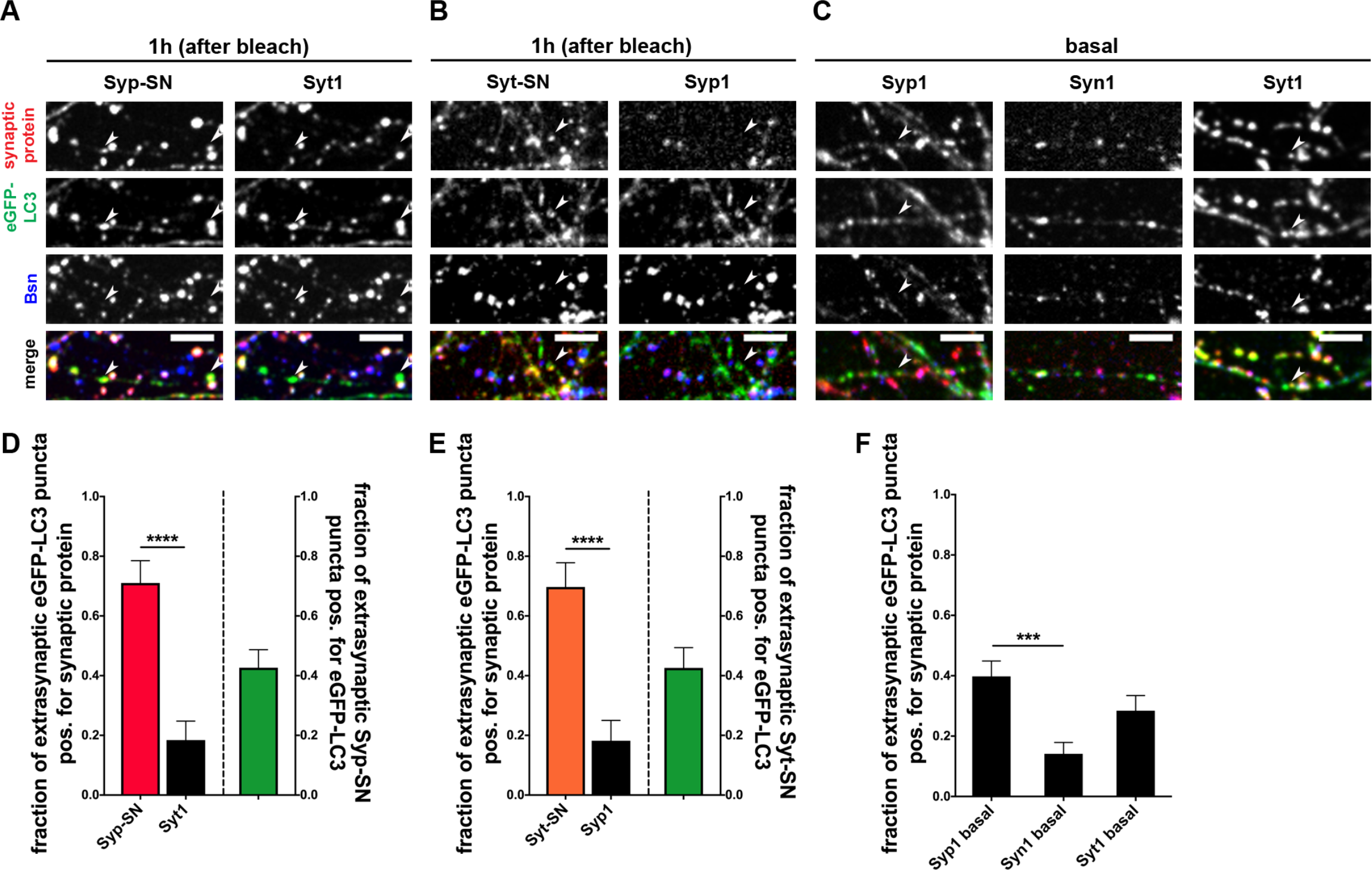
ROS damaged SV proteins selectively accumulate in autophagy organelles. (A and B) Images of hippocampal neurons expressing Supernova-tagged synaptic proteins Syp-SN (A) or Syt-SN (B) that were fixed 1 hour after bleaching and stained with antibodies against GFP, Supernova, Bassoon and Synaptotagmin1 (Syt1) (A) or Synaptophysin1 (Syp1) (B). (C) Images of hippocampal neurons expressing eGFP-LC3 only were fixed untreated (basal autophagy) and stained with antibodies against GFP, Bassoon and Synaptophysin1, (Syp1) or Synapsin1 (Syn1), or Synaptotagmin1 (Syt1). (D) Quantification of the fraction of extrasynaptic eGFP-LC3 puncta positive for SN-tagged Synaptophysin, indicated by arrowheads in (A), or endogenous Syt1 within the same experiment (Syp-SN = 0.71 ± 0.075, n = 38 puncta, 3 independent experiments; Syt1 = 0.18 ± 0.064, n = 38 puncta, 3 independent experiments, p<0.0001). Also quantified is the fraction of extrasynaptic Syp-SN puncta that are positive for eGFP-LC3 (0.43 ± 0.060, n = 68 puncta, 3 independent experiments). (E) Quantification of the fraction of extrasynaptic eGFP-LC3 puncta positive for SN-tagged Synaptotagmin, indicated by arrowheads (B), or endogenous Syp1 within the same experiment (Syt-SN = 0.70 ± 0.081, n = 33 puncta, 2 independent experiments; Syp1 = 0.18 ± 0.068, n = 33 puncta, 2 independent experiments, p<0.0001). Also quantified is the fraction of extrasynaptic Syt-SN puncta that are positive for eGFP-LC3 (0.43 ± 0.068, n = 54 puncta, 2 independent experiments). (F) Quantification of the fraction of extrasynaptic eGFP-LC3 puncta also positive for endogenous Synaptophysin1, Synapsin1 or Synaptotagmin1. Note, the fraction of extrasynaptic eGFP-LC3 puncta positive for Synaptophysin1 is significantly higher than the fraction positive for Synapsin1 (Syp1 basal = 0.40 ± 0.051, n = 93 puncta, 2 independent experiments; Syn1 basal = 0.14 ± 0.038, n = 85 puncta, 2 independent experiments; Syt1 basal = 0.28 ± 0.050, n = 81 puncta, 2 independent experiments). Scale bars: 10μm. Error bars represent SEM. Unpaired T-test (D and E) and ANOVA Tukey’s multiple comparisons test (F) was used to evaluate statistical significance.

To confirm the selectivity of autophagic cargo after Supernova-induced damage, we also quantified the fraction of Syt-SN positive extrasynaptic autophagy cargo organelles 1 hour after bleaching. As with Syp-SN, more than 65% of the extrasynaptic eGFP-LC3 puncta colocalized with Syt-SN, while only 18% of the endogenous Synaptophysin1 (Syp1) was present at these sites (Figure 8B and E). These data indicate that the autophagic machinery within presynaptic boutons can detect and selectively remove damaged SV proteins.

The low but significant presence of endogenous Synaptophysin1 and Synaptotagmin1 in eGFP-LC3 puncta could arise either from the peripheral damage of ROS or be part of the basal flux of these proteins through this pathway. We thus examined whether under basal condition endogenous Synaptophysin1 and Synaptotagmin1 versus Synapsin1 associate with extrasynaptic eGFP-LC3 puncta. Here, we observed that higher levels of both endogenous Synaptophysin1 and Synaptotagmin 1 were found at extrasynaptic eGFP-LC3 puncta compared to Synapsin1 (Figure 6C and F). These data indicate that these SV proteins may be cleared through this degradative pathway.

### Synaptic autophagy acts as a beneficial surveillance mechanism maintaining synapse function

A fundamental question within the synaptic proteostasis field is what roles do different clearance systems play during synaptic transmission. Most studies to date on autophagy rely either on the analysis of genetic ablation and inactivation of key autophagic proteins (Atg5 and Atg7) (Rubinsztein et al., 2011; Russell et al., 2014) or the activation of autophagy with drugs like rapamycin, none of which are specific for the synapse and generally trigger a homeostatic response from other systems masking a specific role of autophagy in the system. Having shown that light-induced ROS production can be used to rapidly (5 minutes) trigger the autophagic clearance of selectively damaged SV proteins, we were keen to explore whether presynaptic autophagy contributes to the real-time maintenance of synaptic function. As an initial test of this concept, we examined whether the ROS-induced damage of Synaptophysin-Supernova and subsequent induction of autophagy affected the functional recycling of SVs based on the activity dependent uptake of the styryl dye FM 1-43 (Cochilla et al., 1999). This was accomplished by performing FM1-43 uptake experiments approximately 5 minutes after photobleaching Syp-SN positive boutons. Interestingly, no difference in the efficiency of FM dye uptake could be detected between bleached and unbleached synapses under basal conditions (Figure 9A and D). These data indicate that either the damage created during ROS production is too gentle to affect synaptic function or that the induced autophagy (Figure 2) is sufficient to remove these damaged SV proteins thus maintaining synaptic function. To test this concept, we added 1μM wortmannin 1 minute before photobleaching and maintained it in the tyrodes buffer before and during loading synapses with FM1-43. Intriguingly, under these conditions, the ROS-induced damage of Syp-SN decreases the subsequent loading of FM1-43 within the bleached area compared to those outside (Figure 9B). We quantified the amount of FM dye uptake dependent on the amount of Syp-SN present at the bouton assuming that more Syp-SN causes more damage to the terminal. ROS mediated damage of Syp-SN had no effect on the slope of the linear regression analysis when autophagy was allowed to operate normally (Figure 9C and D), but significantly reduced the slope when the induction of autophagy was blocked with 1μM wortmannin (Figure 9C and E). Importantly, the addition of wortmannin alone did not alter the uptake of FM1-43 or the slope of the linear regression analysis (Figure 9B, C and E). These data indicate that autophagy may operate in real-time to maintain the integrity and functionality of SVs.

**Figure 9.**
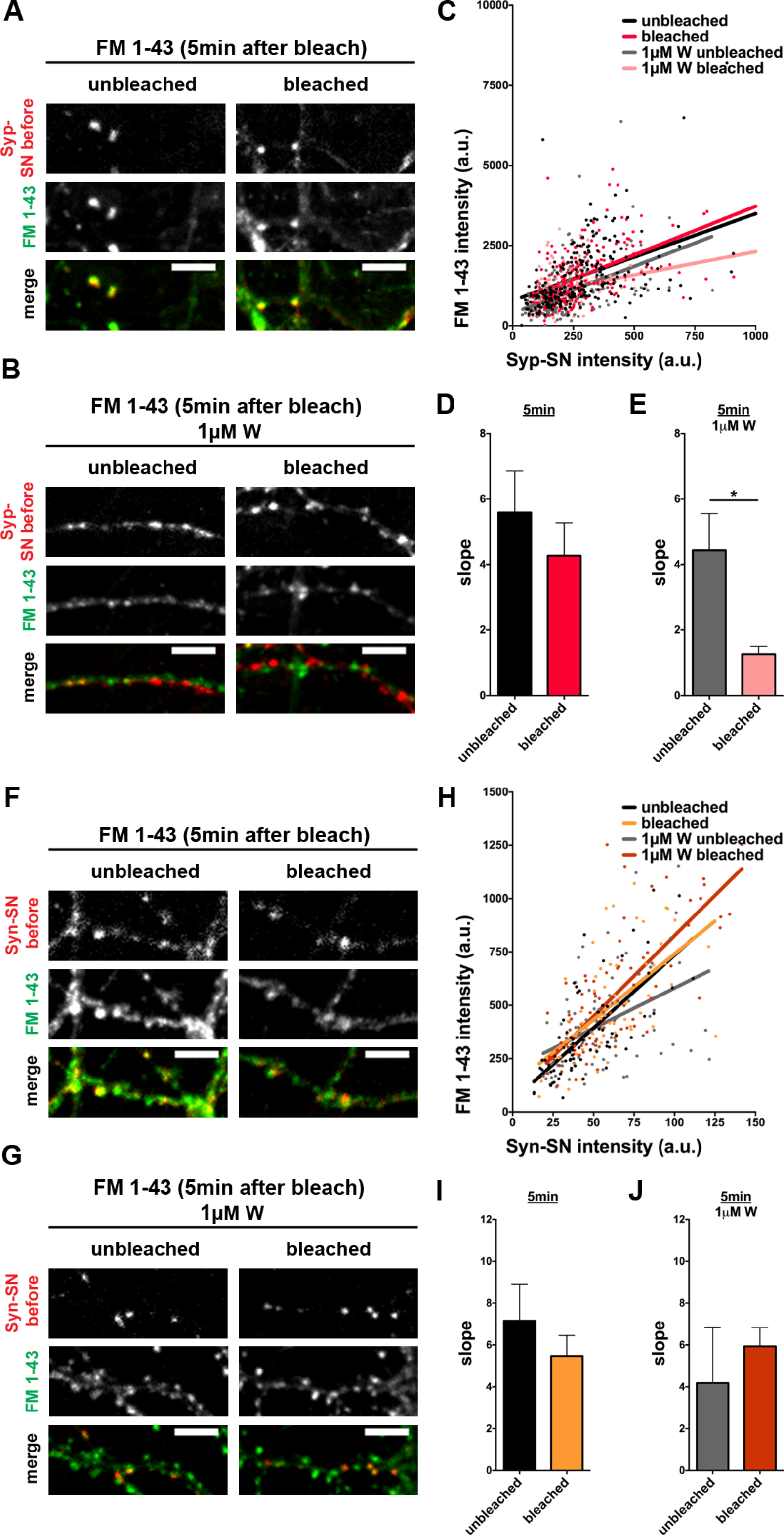
ROS induced damage to Syp-SN impairs FM 1-43 uptake only when autophagy is inhibited. (A, B, F, G) Images of hippocampal neurons expressing FU-Syp-Supernova-P2A-eGFP-LC3 (A and B) or FU-Syn-Supernova-P2A-eGFP-LC3 (F and G), loaded with FM 1-43 for 90 sec in 90mM KCl, 5 min after photobleaching in the absence (A and F) or presence of 1μM wortmannin (B and G). Note, Syp-SN and Syn-SN images were taken before bleaching. (C) Correlation of FM 1-43 intensity over Syp-SN intensity of (A) and (B) (n = 238 (unbleached), 221 (bleached), 221 (1μM W unbleached), 147 (1μM W bleached) synapses, 4 independent experiments). (D and E) Quantification of the slope of the linear regression between bleached and unbleached synapses either in the absence (D: unbleached = 5.59 ± 1.272, 4 independent experiments; bleached = 4.27 ± 1.007, 4 independent experiments) or presence of wortmannin (E: unbleached = 4.43 ± 1.126, 4 independent experiments; bleached = 1.26 ± 0.235, 4 independent experiments, p=0.0332). Note, in the presence of wortmannin, the slope of the linear regression is significantly reduced in bleached synapses compared to unbleached synapses (E). (H) Correlation of FM 1-43 intensity over Syn-SN intensity of (F) and (G) (n = 69 (unbleached), 64 (bleached), 80 (1μM W unbleached), 88 (1μM W bleached) synapses, 3 independent experiments). (I and J) Quantification of the slope of the linear regression between bleached and unbleached synapses either in the absence (I: unbleached = 7.16 ± 1.762, 3 independent experiments; bleached = 5.47 ± 0.986, 3 independent experiments) or presence of wortmannin (J: unbleached = 4.18 ± 2.674, 3 independent experiments; bleached = 5.94 ± 0.900, 3 independent experiments). Note, wortmannin does not affect the uptake of FM 1-43 dye following ROS mediated damage to Syn-SN. Scale bars: 10μm. Error bars represent SEM. Unpaired T-test was used to evaluate statistical significance.

To assess the specificity of this effect, we examined whether the ROS-induced damage of the peripheral SV protein Synapsin also affected the efficiency of FM dye loading. As above, FM1-43 loading was performed ~5 minutes after damaging Syn-SN. Here, we saw no difference in the extent of FM dye uptake between bleached and unbleached synapses under physiological conditions (Figure 9F and I). We also observed no difference in FM dye uptake between bleached and unbleached synapses in the presence of 1μM wortmannin (Figure 9G and J). It should be noted that light damage to Syn-SN expressing boutons does not induce visible synaptic autophagy during the initial 5 minutes following damage (Figure 6K). This suggests that the superoxide generated by Synapsin-SN only modestly damages SVs compared to the integral membrane protein Synaptophysin-SN. Moreover, these data indicate that autophagy plays a minor role in the clearance of Synapsin-SN.

In order to confirm the light-dependent change in FM dye uptake under autophagy inhibition (Figure 9E), we performed electrophysiological experiments. Here, EPSC amplitudes were recorded from autaptic neurons, infected with FU-Syp-SN-P2A-eGFP-LC3 at 2-3 DIV, at 13-18 DIV. Similar to FM dye uptake experiments, bleaching alone did not robustly change the ESPC amplitude (Figure 10A and D) as well as 1μM wortmannin without bleaching (Figure 10B and D). However, under autophagy inhibition with 1μM wortmannin and bleaching, the decrease in EPSC amplitudes was significantly higher (Figure 10C and D). Together these data indicate that autophagy can play a real-time role in the maintenance of synaptic transmission.

**Figure 10.**
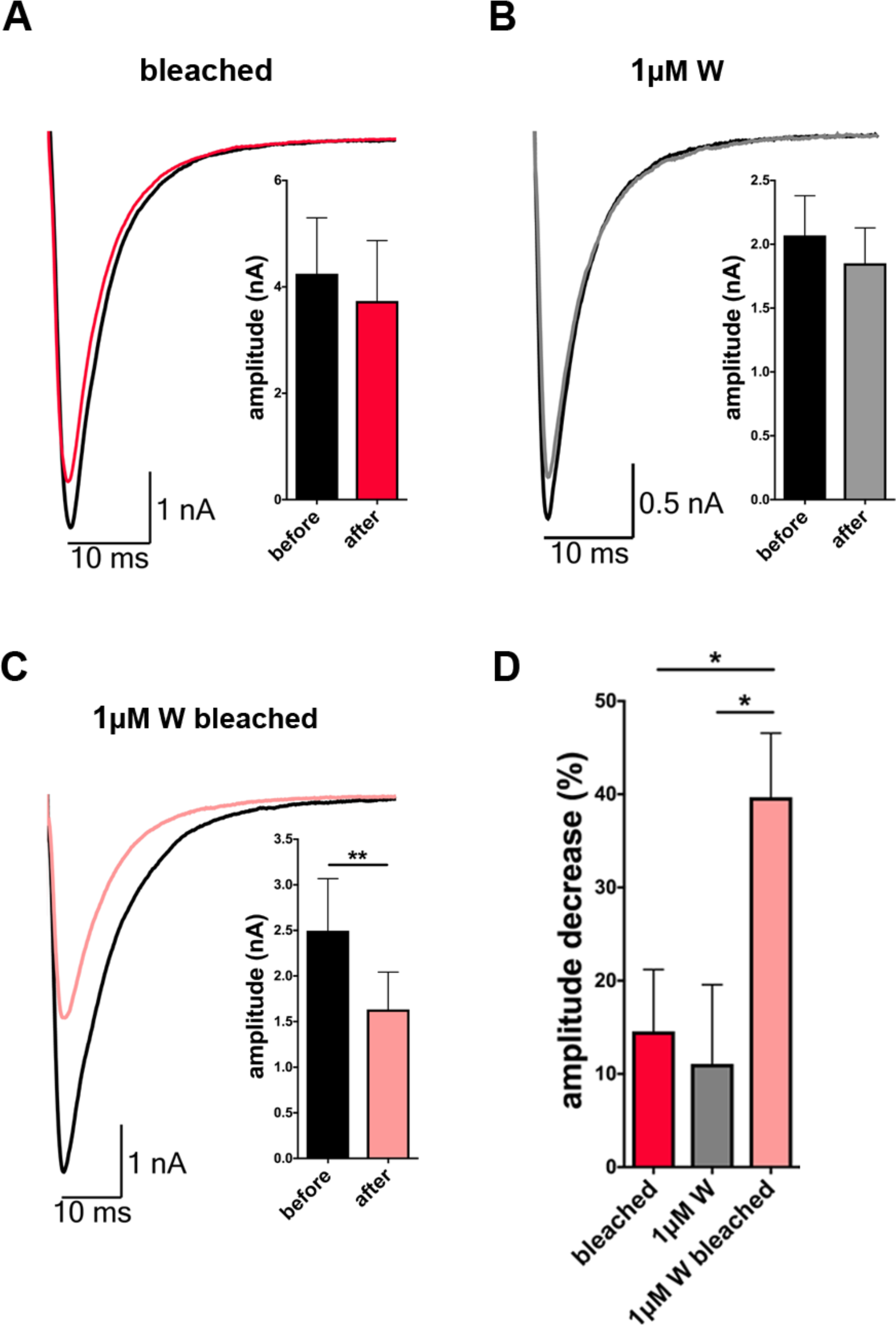
ROS induced damage to Syp-SN impairs evoked release only when autophagy is inhibited. (A, B and C) Example traces of whole-cell patch recording of evoked EPSCs from autaptic hippocampal neurons expressing FU-Syp-Supernova-P2A-eGFP-LC3 before and 5 min after ROS induced damage to Syp-SN either in the absence (A) or presence of 1μM wortmannin (C). Neurons treated with wortmannin but were not bleached served as a control (B). (A amplitude: before = 4.25 ± 1.050, after = 3.74 ± 1.134, 14 neurons, 3 independent experiments) (B amplitude: before = 2.07 ± 0.311, after = 1.85 ± 0.277, 13 neurons, 3 independent experiments) (C amplitude: before = 2.50 ± 0.570, after = 1.63 ± 0.409, 16 neurons, 3 independent experiments, p=0.0042) (M) Quantification of the percent decrease in EPSC amplitude after photobleaching. Note that the decrease in amplitude is significantly higher when 1μM wortmannin is present (bleached = 14.57 ± 6.620, n = 14 neurons, 3 independent experiments; 1μM W = 11.09 ± 8.479, n = 13 neurons, 3 independent experiments; 1μM W bleached = 39.69 ± 6.869, n = 16 neurons, 3 independent experiments, p=0.0445 and p=0.0223). Error bars represent SEM. Paired T-test (A, B and C) and ANOVA Tukey’s multiple comparisons test (D) was used to evaluate statistical significance.

## Discussion

Mechanisms regulating quality control and turnover of synaptic proteins are fundamental to synapse integrity, however, they are not well understood. In this study, we provide evidence that autophagy can be rapidly induced within presynaptic boutons either by the mTOR inhibitor rapamycin or by the selective damage of SV proteins through superoxides. The time range of autophagy induction is consistent with the concept that the machinery is maintained and regulated within presynaptic boutons. Our data also suggest a real-time role for autophagy in maintaining synaptic function, as without it the accumulation of damaged SV proteins can compromise synaptic transmission.

A prerequisite for a real-time functionality for autophagy within presynaptic boutons is its activation on short time scales (seconds/minutes) after insults that damage presynaptic proteins. Studies show that autophagic organelles appear within axons and presynaptic boutons 3-7 hours following the addition of rapamycin (Hernandez et al., 2012), 48 hours after treatment with Sonic Hedgehog (Petralia et al., 2013) and after neuronal activity (Soukup et al., 2016; Wang et al., 2015). Moreover, Bassoon, as well as presynaptic proteins like Rab26 and Endophilin A have been functionally linked to the autophagy machinery (Binotti et al., 2015; Okerlund et al., 2017; Vanhauwaert et al., 2017) of which Atg5, Atg16, LC3 and p62 have been localized to presynaptic boutons (Okerlund et al., 2017). However, as autophagosomes are highly mobile (Cheng et al., 2015b; Maday et al., 2012), it remains unclear whether they arise within boutons or simply accumulate there.

In the current study, we developed a lentiviral vector, expressing a SV protein and the autophagy marker LC3 to monitor autophagic structures in real-time. Similar to earlier studies (Hernandez et al., 2012), we observed low basal autophagy levels within axons (Figure 1F). However, eGFP-LC3 levels increase within 10 minutes within presynaptic boutons and axons following 2μM rapamycin treatment (Figure 1G, K and L), which is much shorter than reported earlier. eGFP-LC3 accumulation was indeed due to elevated autophagy as its increase was blocked by 1μM wortmannin (Figure 1N, O and P) (Codogno et al., 2011; Mizushima et al., 2011).

Although the induction of axonal and presynaptic autophagy is faster than previously recognized, the addition of rapamycin is neither specific for any one neuronal compartment, nor a natural inducer of autophagy (Deng et al., 2017). Therefore we developed a vector system that allows us to generate a spatiotemporally controlled insult within presynaptic boutons. We made use of the fact that free radicals trigger the damage of proteins *in vivo* (Jarvela and Linstedt, 2014) and tagged synaptic proteins with a genetically encoded photosensitizer Supernova (Takemoto et al., 2013). With similar approaches it has earlier been possible to damage mitochondria and induce mitophagy (Ashrafi et al., 2014; Wang et al., 2012; Yang and Yang, 2011).

ROS induced damage of Synaptophysin led to a very rapid induction of autophagy, indicated by the accumulation of eGFP-LC3 within presynaptic boutons within 5 minutes and its spread into axons over time (1-2 hours) (Figure 2I and K). The rapid temporal accumulation of LC3 in boutons was also observed for autophagic vacuoles, as detected by electron microscopy (Figure 4D). These data indicate that the autophagy machinery is present within presynaptic terminals and can be engaged following a local insult within minutes.

A fundamental question raised by our study is whether ROS damage to SV proteins exclusively turns on autophagy as clearance mechanism or multiple protein degradation systems. The endo-lysosomal system has been reported to also clear SV proteins in response to ongoing synaptic activity (Sheehan et al., 2016; Uytterhoeven et al., 2011). One hallmark of this pathway is the appearance of multivesicular bodies (MVB) (Raiborg and Stenmark, 2009). We failed to observe a significant increase in the number of synaptic MVBs on electron microscopy level (Figure 4E and H), indicating that in contrast to autophagy, the endo-lysosomal system is not robustly engaged by ROS mediated protein damage. However, monitoring Rab7 and Chmp2b levels (Sheehan et al., 2016; Stenmark, 2009) by light microscopy, we did observe a modest increase in their colocalization with Synaptophysin-SN, following ROS production (Figure 5). Thus we cannot rule out that ROS mediated damage to SV proteins can trigger the activation of several degradative systems. Interestingly, the study from Sheehan et al. (2016) shows that only a subset of SV proteins is preferentially degraded by the endo-lysosomal system, highlighting the importance to address in the future if distinct SV proteins are degraded via specific and therefore separate pathways and how they are being tagged.

Our data indicate that autophagy induction is not sole dependent on the damage of Synaptophysin. Also the destruction of Synaptotagmin as well as Synapsin leads to elevated autophagy levels at presynapses (Figure 6). However, eGFP-LC3 levels increased with a slower time course (~1 hour time range) (Figure 6). The discrepancy could be indicative for less ROS mediated damage to these SV proteins, possibly due to an increased distance of Supernova from the surface of SVs. Synaptotagmin has a large cytoplasmic region (346aa; comprised of two C2 domains) (Ybe et al., 2000), compared to the shorter 95aa C-terminal tail in Synaptophysin (Gordon and Cousin, 2014) and Synapsin is only peripherally associated with SV membranes, thus further away. An interesting feature of Synapsin is that only about 50% is directly bound to SVs and that it is released from SVs during synaptic activity (Cesca et al., 2010; Chi et al., 2001). This raises the interesting question whether the association of the protein with SVs is necessary to induce autophagy after Synapsin damage. Indeed, dis-engaging Synapsin-SN from SVs during light triggered ROS production did not induce elevated eGFP-LC3 levels (Figure 7E and F). These data support the hypothesis that it is the ROS mediated damage to SVs that triggers the activation of presynaptic autophagy.

An additional question raised by our study is whether the clearance mechanisms triggered by ROS mediated damage leads to the selective removal of only the damaged SV protein or the elimination of the entire associated SV. Indeed, about 70% of extrasynaptic eGFP-LC3 puncta were also positive for Synaptophysin-SN following ROS-mediated damage (Figure 8D). Intriguingly, a much smaller fraction (18%) of these puncta were positive for endogenous SV protein such as Synaptotagmin, which is not known to directly interact with Synaptophysin (Bonanomi et al., 2007; Rizzoli, 2014). These data suggest that presynaptic autophagy can specifically remove damaged SV proteins from synapses. This concept is supported by a reciprocal performed experiment with neurons expressing Synaptotagmin-SN. Here, we also observed a dramatic recruitment of Synaptotagmin-SN into extrasynaptic eGFP-LC3 puncta, but only modest levels of endogenous Synaptophysin (Figure 8E), implying that ROS mediated damage caused by Supernova is rather limited and primarily affects co-tethered proteins, a concept consistent with the low quantum yield of Supernova (Shu et al., 2011; Trewin et al., 2018) and the limited damage half-radius of 3-4nm (Takemoto et al., 2013). Presumably, using the more potent photosensitizer miniSOG (Lin et al., 2013; Qi, 2012; Shu et al., 2011), more SV proteins could be damaged, causing a much bigger insult perhaps leading to the wholesale removal of SVs. Conceptually, a selective removal model also makes metabolic sense, as it would allow for the differential removal of specific mis-folded or damaged proteins, consistent with different half-lives of SV proteins (Cohen et al., 2013).

The rapid induction of presynaptic autophagy within minutes suggests that it possibly has real-time functions at synapses, e.g. helping to maintain synaptic health and integrity. Studies by Lin et al. (2013) showed that targeting miniSOG via Synaptophysin or VAMP2 to SVs leads to a real-time disruption of neurotransmitter release following light activation. Although the precise mechanism was not investigated, loss of function is most likely due to the inactivation of the tagged and/or neighboring SV proteins (Jacobson et al., 2008; Qi, 2012). In contrast, in our experiments with Supernova-tagged Synaptophysin, we did not observe an overt change in synaptic function, assessed by the uptake of FM1-43 dye or the evoked release of neurotransmitter following light-induced ROS production (Figure 9). This suggests that protein damage caused by Supernova radiation is less potent than that induced with miniSOG-Synaptophysin. Intriguingly, when the induction of autophagy was blocked during ROS-mediated damage of Synaptophysin-SN, the extent of FM1-43 uptake, as well as the evoked EPSC amplitude, were reduced (Figure 9 and 10). These data suggest that synaptic autophagy may function in real-time to remove damaged SV proteins contributing to the maintenance of synaptic function. This is consistent with the real-time increase in presynaptic autophagy following ROS mediated damages (Figure 2 and 3).

Taken together, these real-time generators of ROS can represent a powerful tool to spatiotemporally induce damage to synapses and thus increase our understanding of how different clearance systems function during both health and disease. This will be particularly important for the study of neurodegenerative disease where the proper function of autophagy and the endo-lysosmal systems are thought to be crucial for neuronal health (Nixon, 2013; Rubinsztein et al., 2012).

Author contributions: S. Hoffmann preformed the majority of the experiments and analyzed data. S. Hoffmann, F. Ackermann, C. Rosenmund and C.C. Garner designed experiments. M. Orlando performed electron microscopy studies and E. Andrzejak performed electrophysiology experiments. Vectors were generated by S. Hoffmann and T. Trimbuch. S. Hoffmann, F. Ackermann and C.C. Garner wrote the manuscript.

## Acknowledgments

We would like to thank Prof. Eckart D. Gundelfinger and Noam E. Ziv for discussion and valuable comments on the manuscript, Anny Kretschmer and Christine Bruns for technical assistance. The Virus Core Facility of the Charité - Universitätsmedizin Berlin for virus production. The work was supported by Deutsches Zentrum für Neurodegenerative Erkrankungen (DZNE), the Federal Government of Germany (DFG) SFB958 to CCG.

